# Large-scale discovery of neural enhancers for cis-regulation therapies

**DOI:** 10.1101/2025.11.04.686611

**Authors:** Troy A. McDiarmid, Nicholas F. Page, Florence M. Chardon, Riza M. Daza, George T. Chen, Michael Kosicki, Lucas M. James, Hannah C. Nourie, Dianne Laboy-Cintrón, Arthur S. Lee, Paula Vij, Diego Calderon, Jean-Benoît Lalanne, Beth K. Martin, Kyle Fink, Michael E. Talkowski, Alysson R. Muotri, Benjamin D. Philpot, Len A. Pennacchio, Daniel H. Geschwind, Stephan J. Sanders, Nadav Ahituv, Jay Shendure

## Abstract

CRISPR-based gene activation (CRISPRa) has emerged as a promising therapeutic approach for neurodevelopmental disorders (NDD) caused by haploinsufficiency. However, scaling this *cis*-regulatory therapy (CRT) paradigm requires pinpointing which candidate *cis*-regulatory elements (cCREs) are active in human neurons, and which can be targeted with CRISPRa to yield specific and therapeutic levels of target gene upregulation. Here, we combine Massively Parallel Reporter Assays (MPRAs) and a multiplex single cell CRISPRa screen to discover functional human neural enhancers whose CRISPRa targeting yields specific upregulation of NDD risk genes. First, we tested 5,425 candidate neuronal enhancers with MPRA, identifying 2,422 that are active in human neurons. Selected cCREs also displayed specific, autonomous *in vivo* activity in the developing mouse central nervous system. Next, we applied multiplex single-cell CRISPRa screening with 15,643 gRNAs to test all MPRA-prioritized cCREs and 761 promoters of NDD genes in their endogenous genomic contexts. We identified hundreds of promoter- and enhancer-targeting CRISPRa gRNAs that upregulated 200 of the 337 NDD genes in human neurons, including 91 novel enhancer-gene pairs. Finally, we confirmed that several of the CRISPRa gRNAs identified here demonstrated selective and therapeutically relevant upregulation of *SCN2A, CHD8, CTNND2* and *TCF4* when delivered virally to patient cell lines, human cerebral organoids, and a humanized mouse model of *hTcf4*. Our results provide a comprehensive resource of active, target-linked human neural enhancers for NDD genes and corresponding gRNA reagents for CRT development. More broadly, this work advances understanding of neural gene regulation and establishes a generalizable strategy for discovering CRT gRNA candidates across cell types and haploinsufficient disorders.

## Introduction

Neurodevelopmental disorders (NDD) are a heterogeneous group of conditions affecting nervous system development and are among the leading causes of childhood disability worldwide^1,2^. NDDs collectively affect an estimated 17% of children^1,2^, with severe forms impacting 2%^1–3^. Despite their prevalence and burden, effective treatments for NDDs remain limited, representing a major unmet need^4–6^.

The genetic architecture of NDDs includes both common polygenic and rare monogenic components, as well as rare structural variants that may impact multiple genes^7–13^. Notably, a large fraction of severe NDDs arise from *de novo* point mutations or structural variants. Over the past 15 years, exome or genome sequencing of parent-child trios has implicated *de novo* events at hundreds of loci in NDD^8–11^. These studies have also revealed haploinsufficiency—where functional loss of one copy leaves insufficient expression from the other—as the predominant mechanism underlying monogenic NDDs^8–10,14,15^. For example, 87% (89/102) of high-confidence autism spectrum disorder (ASD) risk genes are predicted to be haploinsufficient^8^, and more broadly, variants conferring haploinsufficiency across hundreds of genes are thought to underlie diverse forms of NDD^8–10,14,16–18^.

These observations motivate the exploration of therapies for NDD that directly address haploinsufficiency. Conventional gene replacement strategies, such as viral cDNA delivery, face critical limitations; for example, the packaging capacity of adeno-associated virus (AAV), optimally 4.7 kb^19^, excludes many neuronal genes, and heterologous overexpression often drives non-physiological expression across all transduced cell types. In contrast, cis-regulation therapy (CRT) employs epigenome editing to boost expression of the intact allele, thereby compensating for loss of the other^20,21^. For example, CRISPR activation (CRISPRa) of the *Sim1* promoter rescues obesity in haploinsufficient mice^20,21^.

CRT offers several key advantages: 1) it bypasses AAV packaging constraints that limit cDNA-based therapies^6,19,20^; 2) it acts through endogenous regulatory elements such as enhancers and promoters, enabling physiological, cell type-specific expression rather than uniform overexpression^20–23^; and 3) by using nuclease-deficient genome editors such as CRISPRa, it avoids introducing on-target or off-target mutations. Together, these features position CRT as a promising therapeutic paradigm for haploinsufficient NDDs.

Preclinical studies have begun to support this potential. For example, CRISPRa-based CRT has been shown to rescue disease-relevant phenotypes in models of haploinsufficiency for genes including *SCN1A* (epilepsy), *KCN1A* (epilepsy), *TCF4* (Pitt-Hopkins syndrome, PTHS), *SCN2A* (ASD) and *CHD8* (ASD)^24–28^. However, while these findings highlight the therapeutic promise of CRT, only a handful of NDD risk genes have been subjected to this paradigm to date, underscoring the need for systematic strategies to extend CRT to the hundreds of genes implicated in haploinsufficient NDDs.

A central challenge for CRT is the need for detailed knowledge of gene regulatory circuitry: determining which cis-regulatory elements (cCREs) are active in the relevant neuronal cell types, mapping the target genes they control, and pinpointing which elements can be leveraged to achieve therapeutic levels of upregulation. In addition, it requires identification of reagents (*e.g.*, specific CRISPRa guide RNAs) that reliably confer this activity. To address these challenges, we combined massively parallel reporter assays (MPRAs)^29,30^ and multiplex single cell CRISPRa screening^22,31^ in human neurons to test thousands of candidate cis-regulatory elements (cCREs) predicted to regulate a set of 337 likely haploinsufficient NDD risk genes. We identify 2,422 MPRA-active neural cCREs and demonstrate that several can be targeted with CRISPRa to drive specific and therapeutically meaningful levels of upregulation. These data provide a systematic resource of active, target-linked cCREs for NDD risk genes and establish a foundation for scaling CRT across the heterogeneous genotypic and phenotypic landscape of NDDs. More broadly, this work expands our understanding of the regulatory architecture of human neurons and illustrates how functional genomics can be harnessed to accelerate therapeutic development for genetically defined disorders of the developing brain.

## Results

### Selection of 337 NDD risk genes and 5,425 candidate neuronal enhancers

We first sought to prioritize NDD or neuropsychiatric risk genes with evidence for haploinsufficiency by performing a meta-analysis of 42,320 cases with parental data from the Deciphering Developmental Disorders study^9^, the Autism Sequencing Consortium^8^, and other studies^16–18^ (**Fig. 1a**). Across these datasets, there were 33,422 *de novo* missense variants and 8,251 *de novo* predicted loss-of-function (dnLoF) mutations, with the expected enrichment in cases vs. controls (*de novo* missense Observed/Expected = 1.322, *P* < 0.001^32^; dnLoF Observed/Expected = 2.357, *P* < 0.001; **Fig. 1b**). From 466 genes previously implicated in NDD by these studies, we selected 337 genes where the association was predominantly driven by an excess of heterozygous dnLoF variants (**Fig. 1b**; **Fig. S1a-b**; **Table S1**; **Methods**), a pattern consistent with haploinsufficiency as the underlying mechanism.

**Figure 1.**
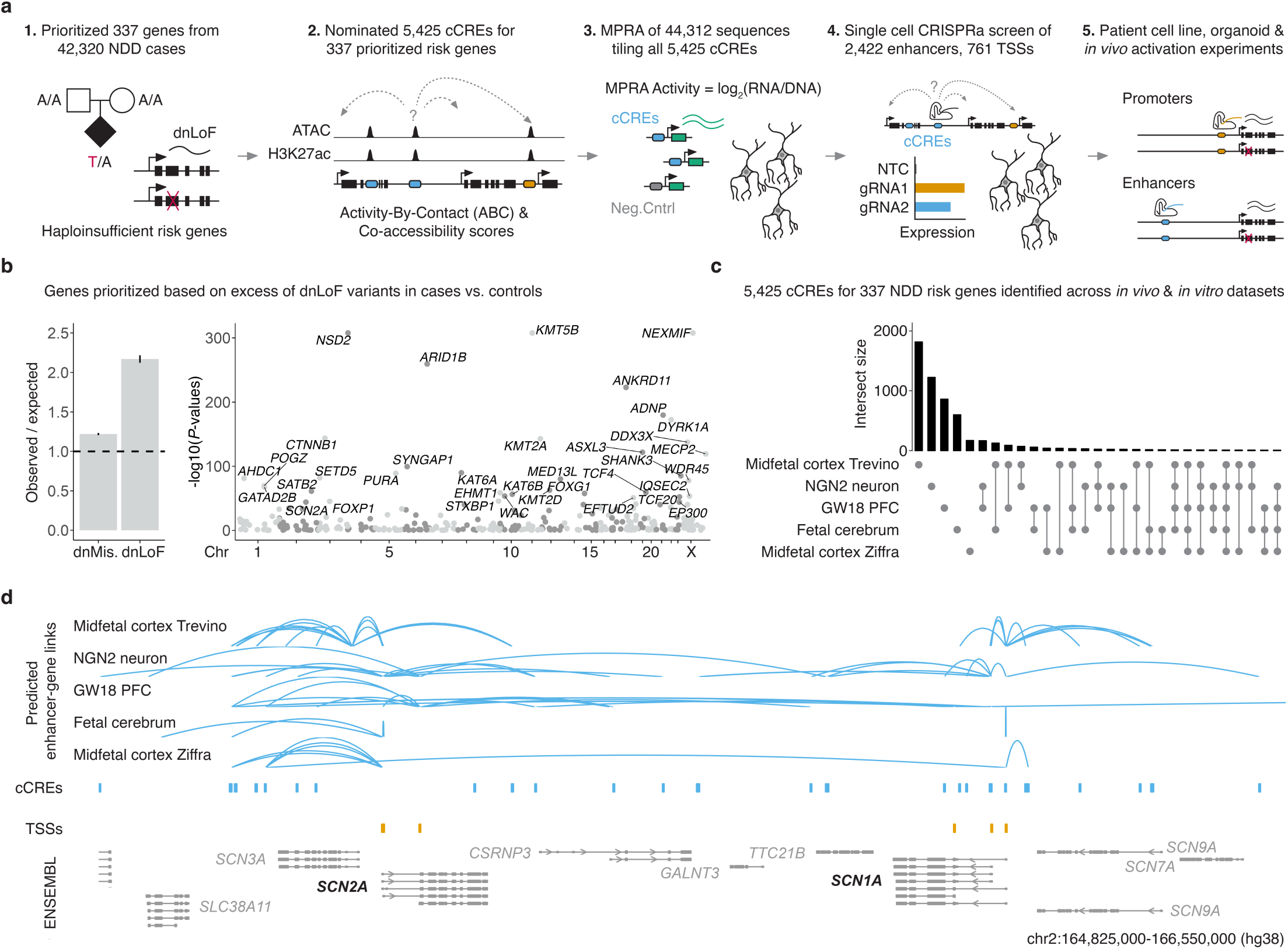
Selection of 337 NDD risk genes and 5,425 candidate neuronal enhancers. **a)** Schematic of workflow. **b)** Risk genes were prioritized based on an excess of dnLoF variants in cases vs. control. (left) Observed/Expected *de novo* missense (dnMis.) and dnLoF variants in cases vs controls. (right) Manhattan plot of observed vs. expected *P*-values for dnLoF mutations. **c)** 5,425 candidate neuronal enhancers, all distal cCREs, for 337 prioritized risk genes were identified using multiple *in vivo* and *in vitro* datasets and enhancer-gene prediction approaches^33,35,38–40^. Upset plot summarizes the sources of these 5,425 cCREs for the 337 prioritized risk genes. **d)** Tracks for predicted cCREs (blue) and linked TSSs (orange) for prioritized NDD risk genes (labeled in black, bottom) are visualized alongside tracks for RefSeq validated transcripts (ENSEMBL/NCBI).

Next, we annotated candidate *cis*-regulatory elements (cCREs) for these 337 NDD risk genes, drawing on five datasets characterizing the regulatory landscape of the developing human brain, together with various enhancer-gene mapping strategies (**Fig. 1c**; **Fig. S1c**; **Methods**). These were: 1) Paired sc-RNA-seq and sc-ATAC-seq data from human midfetal cortex (gestational week [GW] 18-26) and correlation-based mapping^33,34^; 2) Bulk ATAC-seq data from iPSC-derived excitatory neurons (differentiated for 7-8 weeks; NGN2 neurons, as used later in this study) and Activity-By-Contact (ABC) mapping^35,36^; 3) Bulk ATAC-seq data from human midfetal prefrontal cortex (GW18) and ABC mapping^36,37^; 4) Single cell ATAC-seq data from human midfetal cerebral excitatory neurons (GW14-20) and Cicero co-accessibility mapping^38,39^; and 5) Single cell ATAC-seq data from human midfetal cortex excitatory neurons (GW17-21) and ABC mapping^36,40^.

Altogether, we identified 80,533 cCREs across *in vivo* and *in vitro* datasets, including 5,425 candidate enhancers predicted to regulate the aforementioned 337 prioritized NDD risk genes in the developing human brain (**Fig. 1c**; **Fig. S1c**; **Table S2**). To annotate these genes’ promoters, we used capped analysis of gene expression (CAGE) data from the human fetal brain^41^ together with additional annotations^42^ to nominate putative transcription start sites (TSSs), altogether identifying 761 promoters or alternative TSSs (**Fig. 1d**; **Fig. S1k**; **Methods**).

We next compared the 337 prioritized NDD risk genes to all other brain-expressed genes^43^. These NDD risk genes were significantly less tolerant to predicted LoF variation^44^ (*P* < 2.2e-16; **Fig. S1d**), exhibited higher median brain expression^43^ (*P* < 2.2e-16; **Fig. S1e**), and had more candidate neural enhancers on average (*P* < 3.4e-16; **Fig. S1f-i**). These properties further support their functional importance and candidacy for CRISPRa-based CRT.

Interestingly, although 14% (744/5,425) of the candidate neuronal enhancers were predicted by two or more datasets/strategies, most enhancers and predicted enhancer-gene links were identified by one of these (**Fig. 1c-d**; **Fig. S1c,j**; **Table S2**), and correlations between datasets/strategies were generally low (**Fig. S1j**; rho = 0.002-0.18). Taken together, these analyses highlight the limitations of current methods for predicting enhancer–gene links from correlative biochemical data alone, and the need for large-scale functional experiments to resolve which candidate enhancers are truly active in human neurons as well as which gene(s) they regulate.

### MPRAs identify functional neuronal CREs

To determine which of the 5,425 candidate enhancers are active in human neurons, we used a lentivirus-based MPRA (lentiMPRA)^30,45,46^ (**Fig. 2a**). First, we designed, synthesized and cloned an MPRA library consisting of 44,312 x 270 base pair (bp) sequences densely tiling all 5,425 cCREs with 90 bp overlaps (**Methods**). We also included 100 active and 99 inactive control sequences from an MPRA of open chromatin regions conducted in neural precursor cells (NPCs)^47^, 729 active and 600 inactive control sequences from an MPRA of human accelerated regions (HARs) conducted in an SH-SY5Y immortalized neuroblastoma cell line^48^, and 500 base-level shuffled versions of a randomly selected subset of the 44,312 tiles. Altogether, the lentiMPRA library included 46,340 unique designs (**Table S3**), and was cloned such that each design was paired with multiple barcodes residing in the 5’ UTR of the downstream reporter (**Fig. S2a**).

**Figure 2.**
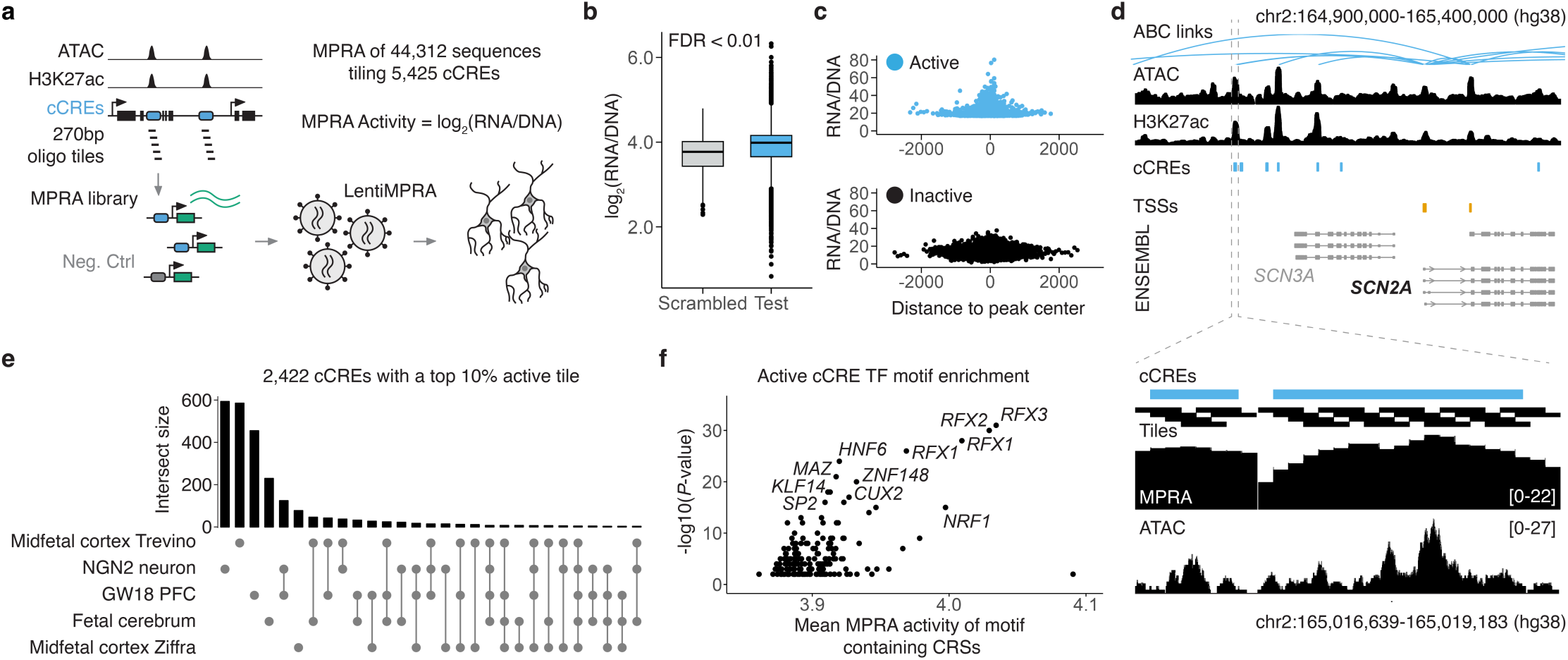
Massively parallel reporter assay (MPRA) identifies cCREs active in human neurons. **a)** Schematic of lentiMPRA experiment. 44,312 x 270-bp oligo sequences tiling 5,425 cCREs were synthesized, cloned to a 5’ lentiMPRA vector, and associated with degenerate 5’ UTR barcodes. This lentiviral MPRA library was packaged and delivered to human iPSC-derived neurons. Integrated DNA and transcribed RNA barcodes were sequenced to quantify the transcriptional activity of each tile. **b)** Boxplot of log2 scaled MPRA activities (RNA/DNA barcode ratios) of tiles derived from candidate cCREs (“test”) vs. scrambled controls (“scrambled”). Boxes represent the 25th, 50th, and 75th percentiles. Whiskers extend from hinge to 1.5 times the interquartile range. Data beyond the whiskers are plotted as individual points. **c)** Scatterplots of MPRA activities (y-axes) of active test tiles (top; FDR < 0.01) or inactive test tiles (bottom; FDR > 0.01) vs. their distance from the center of the ATAC-seq peak from which they were derived (x-axes). **d)** Browser plot of the *SCN2A* locus, together with cCREs that are transcriptionally active in human iPSC-derived neurons. (Top) Tracks for predicted cCREs (blue) and linked TSSs (orange) for prioritized NDD risk genes (labeled in black) are visualized alongside tracks for iPSC-derived neuron ATAC-seq^35^, H3K27ac^35^, and RefSeq validated transcripts (ENSEMBL/NCBI). (Zoom in, bottom) Average MPRA activity scores of test sequences (black bars) densely tiling cCREs (blue bars). Below that, average MPRA activity of cCRE tiles is shown above iPSC-derived neuron ATAC-seq track. **e)** Upset plot summarizes the sources of the 2,422 cCREs that were associated with a tile whose MPRA activity was in the top 10%. **f)** cCRE-derived tiles containing binding motifs of neuronal TFs exhibit significantly higher MPRA activity.

Following stable transduction at high multiplicity of integration (MOI) (mean 46 across 3 replicates) in human iPSC-derived neurons^49,50^, we amplified and sequenced RNA and DNA barcodes to quantify the relative activities of tiled cCREs (**Fig. 2a**). We recovered a median of 163 DNA barcodes per element with 97% (44,706/46,340) associated with at least one barcode (**Fig. S2b**; **Table S3**). MPRA activity scores were well correlated across transduction replicates (*r* = 0.77-0.78; *rho* = 0.76-0.77; **Fig. S2c-e**). On average, candidate enhancer tiles were only slightly more active than scrambled controls (1.05-fold; **Fig. S2f**), but there were 9,547 with significantly greater activity (**Fig. 2b**; FDR < 0.01), and the top 10% exhibited 1.16- to 1.72-fold greater activity than the mean scrambled control. Interestingly, although controls selected for activity in earlier studies were more active than our base-shuffled negative controls (1.11-fold for NPC-active fragments^47^; 1.06-fold for SH-SY5Y-active fragments^48^), so were controls selected for inactivity in these same studies (1.07-fold for NPC-inactive fragments^47^; 1.05-fold for SH-SY5Y-active fragments^48^), potentially because some of these candidate cCREs are more active in differentiated neurons (**Fig. S2f**).

We next analyzed which biochemical marks were associated with active regulatory activity. Active tiles were clustered near the center of chromatin accessibility peaks (**Fig. 2c-d**). MPRA activity scores were also positively correlated with biochemical features associated with enhancer activity in neurons (*e.g.* H3K27ac, H3K4me1, ASCL1 ChIP-seq) and negatively correlated with repressive marks (*e.g.* H3K4me3) (**Fig. S2g**). Active tiles were also enriched for the motifs of transcription factors (TFs) active in human neurons, including *RFX1*, *RFX2*, and *RFX3*^10,51,52^, as well as Universal Stripe Factor TFs^53^, which we have previously observed to be enriched in active neuronal enhancers via lentiMPRA^54^ (**Fig. 2f**).

Of the 5,425 candidate enhancers tiled in this MPRA, 145 were previously tested *in vivo* by means of a mouse enhancer assay^54,55^. Of these, 66% (96/145) drove reporter expression in the developing mouse CNS, and within that subset, 63% (60/96) corresponded to a highly active tile in our MPRA (top 10% of scores; **Fig. S2h**). Conversely, the candidate enhancers that were the source of highly active MPRA tiles were much more likely to be active in the developing mouse CNS (odds ratios: 2.3-3.9; **Fig. S2h**). As further validation, we selected two candidate enhancers that overlapped with highly active MPRA tiles and were predicted to regulate the well-established NDD genes, *CHD2* and *AUTS2*, and subjected them to a similar mouse enhancer assay (**Methods**). Both cCREs were active in the developing mouse CNS (**Fig. S2i-j**), reinforcing the suitability of this approach for prioritizing functional neuronal CREs.

Altogether, 45% (2,422/5,425) of the candidate enhancers overlapped with one or more highly active MPRA tiles (top 10% of scores; **Fig. 2e**; **Table S3**). Although most of these functionally validated 2,422 cCREs were nominated by only a single dataset/strategy (**Fig. 2e**), validation rates were higher for those predicted by multiple methods (1+: 45%; 2+: 63%; 3+: 77%; 4+: 79%). These 2,422 candidate enhancers are collectively predicted to regulate 92% (309/337) of our prioritized NDD risk genes, and the vast majority linked with only one of these genes (**Fig. S2k**). Taken together, our MPRA provided functional support for thousands of biochemically nominated cCREs near NDD risk genes, setting the stage for CRISPRa-based targeting of their regulatory activity in endogenous genomic contexts.

### CRISPRa screen identifies functional CREs for 200 NDD risk genes

We next sought to leverage a multiplex single cell CRISPRa screening framework^22^ to determine which endogenous cCREs could be targeted with CRISPRa to yield upregulation of NDD risk genes in a relevant model of mature human neurons (**Fig. 3a**). First, we designed four gRNAs for each of the 2,422 candidate enhancers prioritized by MPRA, as well as 761 promoters/TSSs of the 337 prioritized NDD risk genes (**Methods**). We supplemented these designs with additional TSS-targeting gRNAs selected from a genome-wide CRISPRa library^56^ and non-targeting control (NTC) gRNAs. Altogether, our library design included 15,643 gRNAs, including 9,685 candidate enhancer-targeting gRNAs, 4,458 TSS-targeting gRNAs, and 1,500 NTC gRNAs (**Fig. S3a**; **Table S4**).

**Figure 3.**
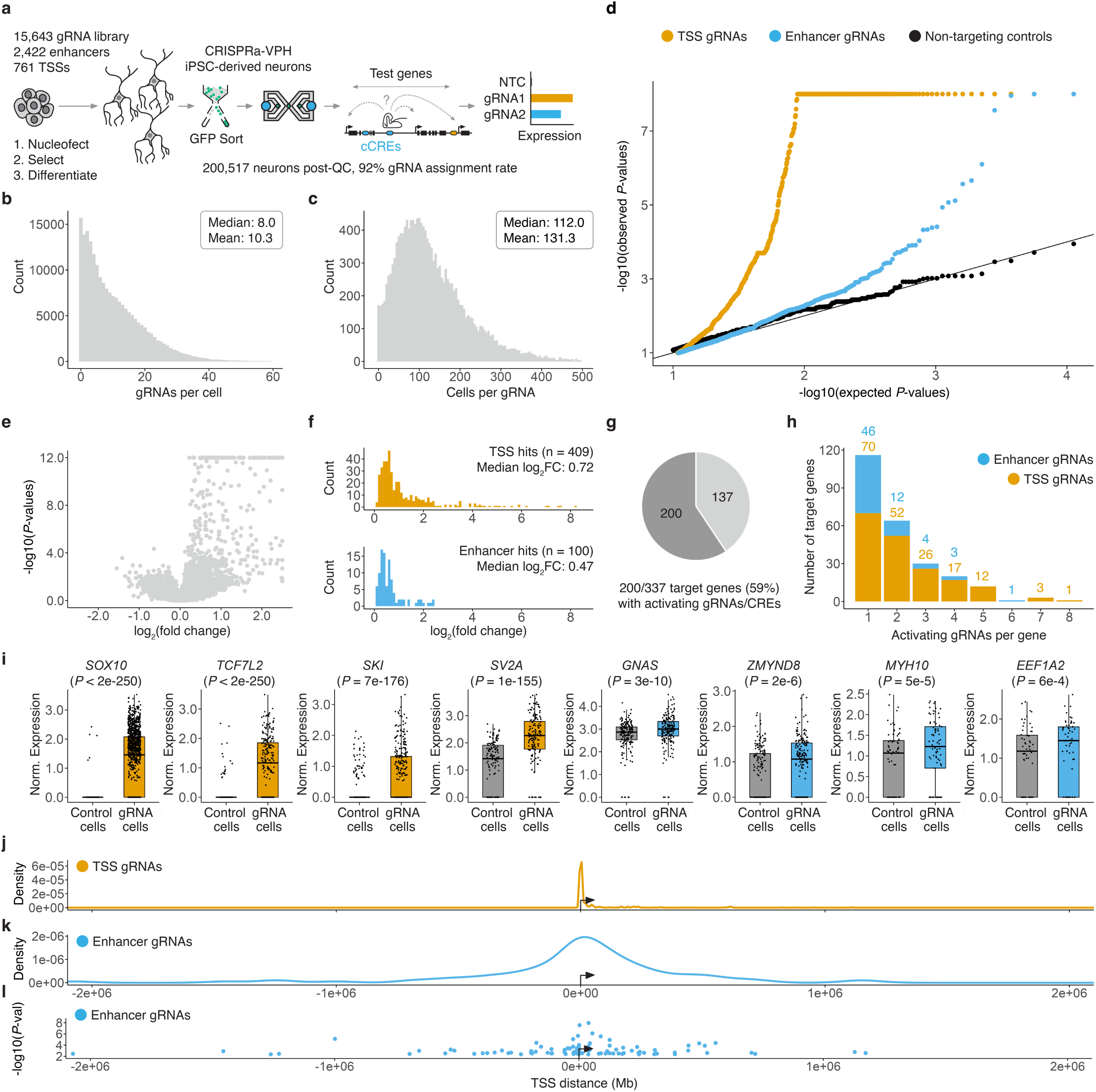
Multiplex, single cell CRISPRa screen identifies functional endogenous cCREs for 200 NDD risk genes. **a)** Schematic of multiplex, single cell CRISPRa screen. **b)** Histogram of number of cells per gRNA following single cell transcriptome QC and gRNA assignment. **c)** Histogram of number of gRNAs detected per cell. **d)** Quantile-quantile plot showing distribution of expected vs. observed *P*-values for promoter-targeting (orange), enhancer-targeting (blue), and non-targeting (black, downsampled) differential expression tests. **e)** Volcano plot showing the average log2 fold-change and *P*-values for target genes. **f)** Histogram of average log2 fold change in expression for activating gRNAs targeting promoters (orange, top) or enhancers (blue, bottom). **g)** Proportion of NDD risk genes for which an activating gRNA was identified. **h)** The number of promoter- and enhancer-targeting gRNAs identified per gene. **i)** Example boxplots for select hit gRNAs showing the log2 fold change in expression between gRNA cells and control cells. Dots represent normalized expression for individual cells. Control cells are downsampled to have the same number of cells as the indicated targeting gRNA for visualization. Boxes represent the 25th, 50th, and 75th percentiles. Whiskers extend from hinge to 1.5 times the interquartile range. **j-l)** Metagene plots showing the distance of an activating gRNA and linked target gene.

We synthesized and cloned these designs into piggyFlex^22,57^, a piggyBac transposon-based gRNA delivery construct with dual fluorophore and antibiotic selection cassettes to enable enrichment of cells bearing integrated gRNAs (**Fig. S3b**). We then stably integrated the library into iPSCs bearing conditional CRISPRa machinery and an inducible NGN2 transcription factor^58^ poised to drive neural differentiation (**Fig. 3a**; **Fig. S3c**). Following antibiotic selection, iPSCs were strongly GFP-positive, consistent with successful integration (**Fig. S3d**). Furthermore, neurons induced from these iPSCs displayed clear morphological differentiation (**Fig. S3e-g**), and we confirmed effective CRISPRa activity in mature neurons using a minP-tdTomato reporter assay^59^ (**Fig. S3e-f**).

Next, we generated single cell transcriptomes for 200,513 human iPSC-derived neurons with direct capture of gRNA transcripts (10x Genomics HT chemistry; **Fig. 3a**). Analysis of the resulting scRNA-seq profiles further confirmed effective neural differentiation, as neural marker genes (*e.g.*, *MAPT*, *NCAM1*, *MAP2*) were well expressed (**Fig. S3h-k**). Following QC, we obtained a 92% gRNA assignment rate (*i.e.* scRNA-seq profiles with at least one assigned gRNA), with a mean of 10.3 distinct gRNAs assigned per cell and 131 cells per gRNA (**Fig. 3b-c**). With this MOI, we estimate that we have statistical power equivalent to profiling >2 million cells under the more conventional “one gRNA per cell” framework.

For each gRNA, we computationally partitioned cells into those with a given gRNA vs. those without, and tested for upregulation of that gRNA’s putative target gene(s). Applying SCEPTRE^60^, we observed a clear excess of significant differential expression test *P*-values for both TSS- and enhancer-targeting gRNAs compared to NTCs (**Fig. 3d**; **Fig. S3l-n**). Significant effects were strongly enriched for upregulation of our intended target genes (**Fig. 3e**; *P* < 2.2e-16). Statistical signal (**Fig. 3d**) and the degree of upregulation (**Fig. 3f**) was generally stronger for promoters/TSSs than enhancers. Importantly, while a few TSS-targeting gRNAs yielded large upregulation levels, most upregulations were modest (**Fig. 3f**; median 1.6 fold-change for promoter-activating gRNAs; median 1.4 fold-change for enhancer-activating gRNAs; **Fig. 3f**; **Table S5**). This is consistent with the observation that the levels of activation achieved by CRISPRa are more constrained than heterologous overexpression^20,22,23,26^.

Altogether, we identified 509 activating gRNAs yielding upregulation of 200/337 (59%) prioritized NDD risk genes (SCEPTRE FDR < 0.1; **Fig. 3g**; **Table S6**). These included 409 activating promoter-targeting gRNAs and 100 enhancer-targeting gRNAs encompassing 91 neural enhancer-gene pairs (**Fig. 3h**; **Table S6**). As with the results of the MPRA, validation rates were higher for enhancer-gene pairs nominated by multiple methods (1+: 3.8%; 2+: 6.0%; 3+: 9.6%; 4+: 18.5%). We identified activating promoter-targeting gRNAs for 181 genes and enhancer-targeting gRNAs for 66 genes (**Table S6**). CRISPRa signal was highly specific—the vast majority of these gRNAs (86%) yielded upregulation of only the predicted target gene and no other genes within 1 Mb.

Altogether, we identified a gRNA yielding specific upregulation for 190/200 (95%) NDD risk genes (**Table S6**; **Fig. S3o**). Many cCREs were successfully targeted by more than one gRNAs (116/297 (39%) of successfully targeted cCREs, including 7/91 enhancers and 109/206 TSSs; **Table S6**). Many of our target genes also had several promoter- or enhancer-targeting gRNAs yielding significant upregulation (138/200 (69%) NDD risk genes with 2+ activating gRNAs; **Fig. 3h**; **Table S6**). This set included genes that were lowly expressed at baseline, as well as genes that were already highly expressed (**Fig. 3i**; **Fig. S3p-q**). Furthermore, while the full set of 337 NDD risk genes were more highly expressed than other genes (**Fig. S1d**; **Fig. S3p**), the 200/337 genes for which we identified activating gRNAs exhibited modestly higher baseline expression than the 137/337 genes for which we did not (1.7-fold higher baseline expression, *P* = 0.0011; **Fig. S3p**). We also observed that successfully targeted genes with lower baseline expression exhibited greater fold-change upregulation (**Fig. S3q**).

Although CRISPRa signal for enhancers clustered near regulated TSSs, only 37/100 enhancer-activating gRNAs targeted sites within 100 kb of their target gene, and there were many examples of successful CRISPRa targeting of enhancers acting over long distances (**Fig. 3k-l**; **Table S6**). For example, we identified three distal enhancers located 392 kb, 599 kb and 622 kb downstream of the gene body that, when targeted, skip several intervening genes to activate *GNAS* (**Fig. S4a**). This example also illustrates how this framework can be leveraged to identify CRISPRa-responsive promoters, as although several *GNAS* TSSs were targeted, all 3 successfully promoter-activating gRNAs targeted the same one of these (**Fig. S4a**). Similarly, for the transcription factor *ZFHX4*^9,61^, we identified five promoter-activating gRNAs that all targeted a single CRISPRa-responsive TSS, as well as several proximal and distal enhancer activating gRNAs (**Fig. S4b**). Taken together, these results reveal neuronal gene regulatory circuitry across multiple loci, and provide a validated set of MPRA-active, CRISPRa-responsive human neural cCREs, together with validated gRNA targeting reagents, for 200 haploinsufficient NDD risk genes.

### CRISPRa-responsive *SCN2A* cCREs drive gene activation in mice and human cerebral organoids

*SCN2A* encodes a neuronal voltage-gated sodium channel, and haploinsufficiency of *SCN2A* is a major risk factor for ASD^10,62^. In the CRISPRa screen, we identified multiple gRNAs that upregulated *SCN2A*, either by targeting its promoter (gRNA 11993: 1.4-fold increase; gRNA 13779: 1.3-fold increase) or an enhancer located 177 kb upstream of its TSS (gRNA 5432: 1.2-fold increase) (**Fig. 4a**; **Table S6**). Tiles spanning this *SCN2A* enhancer also exhibited strong activity in our human neuronal MPRA (**Fig. 4b**). For these promoter- and enhancer-targeting gRNAs, upregulation was specific to *SCN2A*, as neither *SCN1A* (another NDD gene located downstream of *SCN2A*), *SCN3A* (upstream, intervening the targeted *SCN2A* enhancer and TSS), nor any other genes within 1 Mb, exhibited upregulation (**Fig. 4a**; **Table S6**). We also tested the upstream (−177 kb) *SCN2A* enhancer *in vivo* and found it drives tissue-specific activity in the developing mouse midbrain, hindbrain and neural tube at embryonic (E) day 12.5 (**Fig. 4c**; **Fig. S5a**).

**Figure 4.**
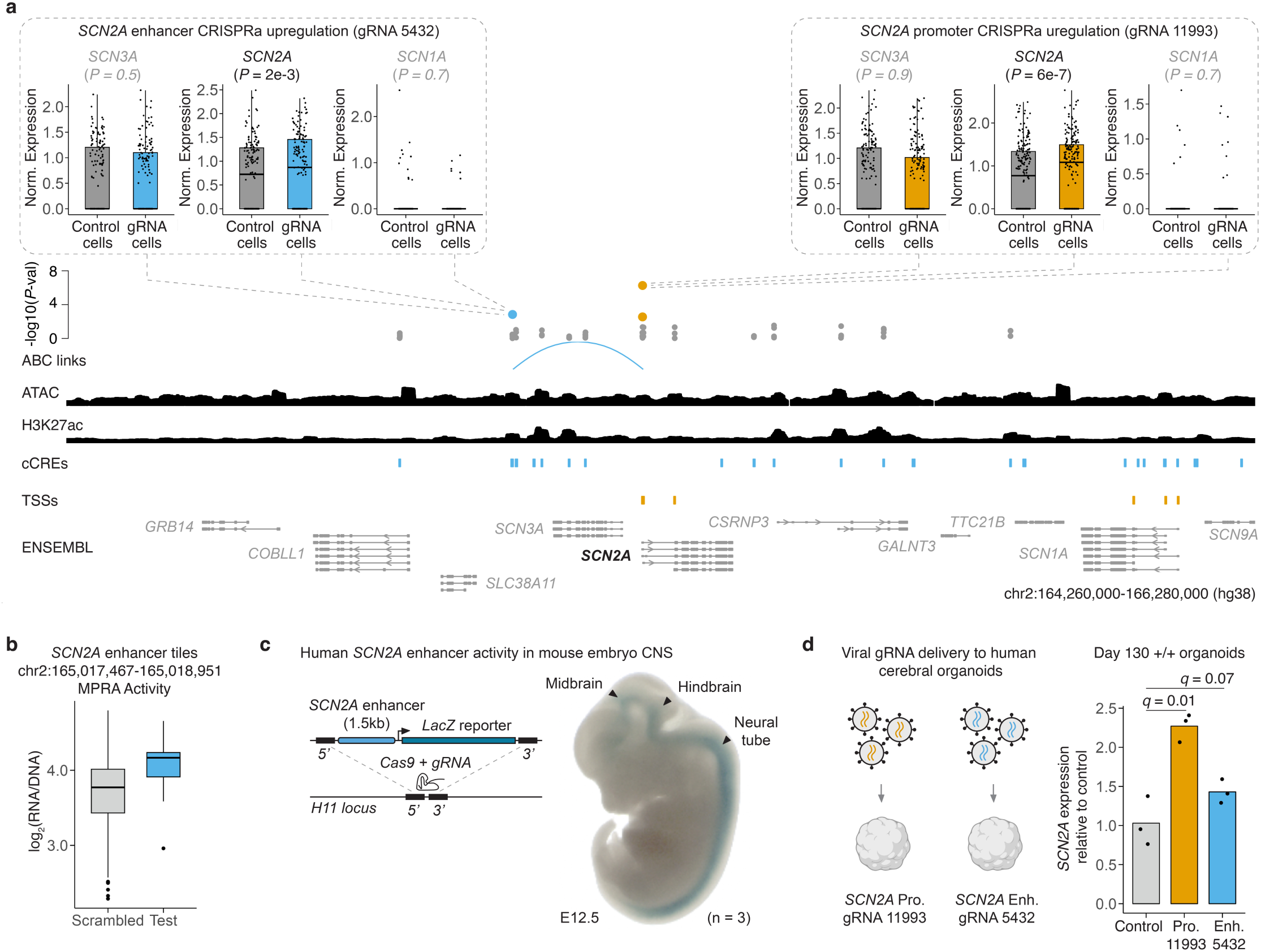
CRISPRa-responsive *SCN2A* cCREs drive gene activation in mice and human cerebral organoids. **a)** (Top) Boxplots showing log2 fold-changes in expression for cells in which specific gRNAs were detected compared to control cells. Example boxplots for *SCN2A* (intended target), and two other genes in the *cis*-neighborhood, *SCN1A* and *SCN3A,* for the indicated enhancer-targeting (boxplots at top left) and TSS-targeting (boxplots at top right) gRNAs are shown. Dots represent normalized expression from individual single cell transcriptomes. Control cells are downsampled to have the same number of cells as the indicated targeting gRNA for visualization. (Bottom) Differential expression test *P*-values for individual *SCN2A* TSS- and enhancer-targeting gRNAs are plotted above tracks for predicted cCREs (blue) and linked TSSs (orange) for prioritized NDD risk genes. TSS- and enhancer-targeting gRNAs yielding significant target gene upregulation (FDR < 0.1) are coloured orange or blue, respectively. These *P-*values are plotted above tracks for iPSC-derived neuron ATAC-seq^35^, H3K27ac^35^, and RefSeq validated transcripts (ENSEMBL/NCBI). Only predicted enhancer gene links that were supported by both MPRA and CRISPRa screen data are shown. **b)** MPRA activity of sequences tiling an enhancer that was successfully targeted in the CRISPRa screen. Coordinates: hg38. **a-b)** Boxes represent the 25th, 50th, and 75th percentiles. Whiskers extend from hinge to 1.5 times the interquartile range. Data beyond the whiskers are plotted as individual points. **c)** (Left) Reporter construct knock-in strategy to test the human *SCN2A* enhancer for activity in mice. (Right) Reporter expression in the CNS of an E12.5 mouse embryo. **d)** (Left) Human cerebral organoids were differentiated for 100 days then transduced with lentiviral constructs encoding either the *SCN2A* promoter-targeting gRNA 11993 or the *SCN2A* enhancer-targeting gRNA 5432. One month after transduction (*i.e.* Day 130 of differentiation), organoids were harvested and *SCN2A* expression was measured using qRT-PCR. (Right) Bars represent average fold-change in *SCN2A* expression (qRT-PCR) relative to empty gRNA vector transduction controls for individual transduction replicates.

Viral vector–mediated delivery is a promising route for implementing CRT^20,21^. To evaluate the therapeutic potential of the *SCN2A*-targeting gRNAs identified in our multiplex CRISPRa screen, we recloned these gRNAs into lentiviral expression constructs. When delivered to HEK293T cells, an easy-to-transduce immortalized cancer cell line with a relatively permissive chromatin environment^63,64^, both promoter- and enhancer-targeting gRNAs upregulated *SCN2A* (**Fig. S5b**; **Table S7**). We next tested whether viral delivery could drive *SCN2A* upregulation in human cerebral organoid cultures^65,66^. Using a stem cell line constitutively expressing dCas9– p300-Core^67^, we established cerebral organoid cultures that we differentiated for 100 days—a developmental stage at which *SCN2A* expression is normally upregulated^68^. Following differentiation, we transduced organoids with either promoter-targeting gRNA 11993 or enhancer-targeting gRNA 5432. One month later, quantitative RT-PCR (qRT-PCR) revealed upregulation of *SCN2A* with either the promoter-targeting (2.3-fold; *q =* 0.01) or enhancer-targeting (1.4-fold; *q =* 0.07) gRNA (**Fig. 4d**).

Notably, the top *SCN2A* promoter-targeting gRNA from our CRISPRa screen, while recognizing an *Streptococcus pyogenes* (Sp) PAM, overlaps an *Staphylococcus aureus* (Sa) gRNA that we previously showed can rescue electrophysiological and behavioral phenotypes in *SCN2A*-haploinsufficient human neurons^28^ (**Fig. S5c**). This convergence underscores the therapeutic relevance of the *SCN2A* cCREs identified here and validates our discovery framework. More broadly, beyond *SCN2A*, our screen revealed hundreds of MPRA-active, CRISPRa-responsive cCREs linked to 200 NDD risk genes, together with gRNAs to which they are responsive in neurons, providing a systematic foundation for the development of CRTs across a wide range of haploinsufficient NDDs.

### Upregulation of NDD risk genes in haploinsufficient cell lines, organoids and mice

We next sought to confirm that CRISPRa gRNAs identified through a multiplex screening framework could be used to upregulate NDD risk genes in diverse models of haploinsufficiency. For this, we used viral vectors to target cCREs for *CHD8*, *CTNND2*, and *TCF4* in haploinsufficient patient-derived cell lines, humanized mice, and/or human cerebral organoids.

*CHD8* encodes a chromatin remodeler that regulates numerous genes important for neurodevelopment, including other NDD risk genes^69–71^. Variants resulting in *CHD8* haploinsufficiency cause a genetic syndrome characterized by increased risk for ASD, ID, and macrocephaly^8,9,69,71^. In the multiplex CRISPRa screen, we identified three activating gRNAs targeting two alternate TSSs that drove upregulation of *CHD8* in human neurons (**Fig. 5a**; **Table S6**). We focused on the gRNA targeting the TSS of the full-length *CHD8* isoform and tested its ability to upregulate *CHD8*. After initial optimization in HEK293T cells (**Fig. 5b**), we established cerebral organoid cultures from *CHD8* heterozygous human stem cell lines. We transduced these organoids with gRNA-12941 after 25 days of differentiation, the developmental timepoint in which *CHD8* is most highly expressed in this culture system^68^. Upon harvesting the organoids cultures at 30 days post-transduction and performing qRT-PCR, we observed upregulation of *CHD8* upon CRISPRa targeting of the TSS of the full-length isoform (2.3-fold; *q =* 0.003; **Fig. 5b**).

**Figure 5.**
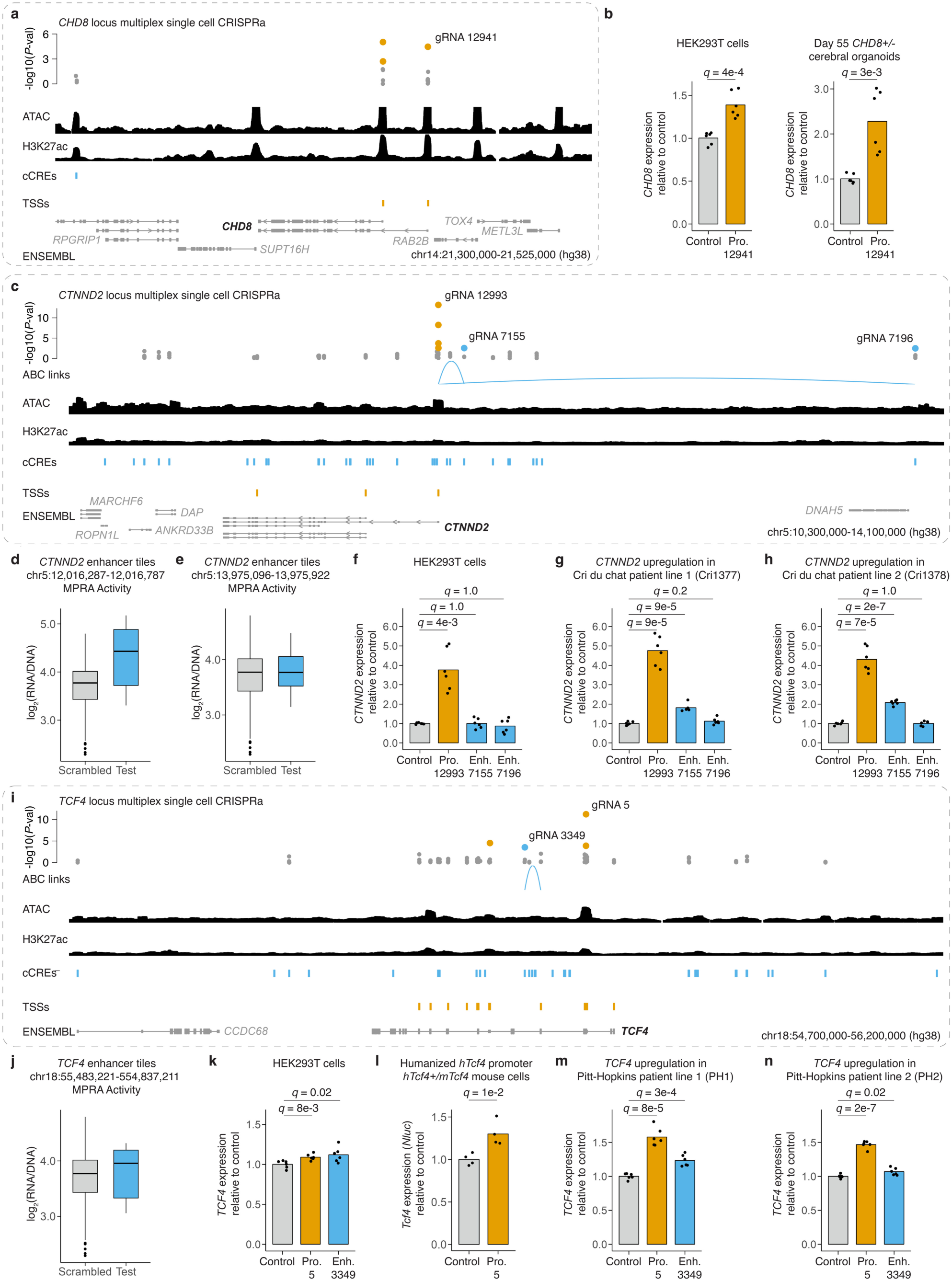
cCREs for *CHD8*, *CTNND2*, and *TCF4* yield upregulation in patient cell lines, humanized mice, and human cerebral organoids. **a)** Multiplex single cell CRISPRa screen differential expression test *P*-values for individual *CHD8* TSS- and enhancer-targeting gRNAs are plotted above tracks for predicted cCREs (blue) and linked TSSs (orange) for prioritized NDD risk genes. TSS- and enhancer-targeting gRNAs yielding significant target gene upregulation (FDR < 0.1) are coloured orange or blue, respectively. gRNAs used in viral validations are labeled. Results are plotted above tracks for iPSC-derived neuron ATAC-seq^35^, H3K27ac^35^, and RefSeq validated transcripts (ENSEMBL/NCBI). Predicted enhancer gene links that were supported by both MPRA and multiplex single cell CRISPRa screens are shown. **b)** Viral delivery of a gRNA targeting a *CHD8* TSS in HEK293T cells (left) and *CHD8+/-* human cerebral organoids (right). Bars represent average fold change in *CHD8* expression (qRT-PCR) relative to empty gRNA vector transduction controls for individual transduction replicates. **c)** Multiplex single cell CRISPRa screen differential expression test *P*-values for individual *CTNND2* TSS- and enhancer-targeting gRNAs. Tracks as in panel **a**. **d-e)** MPRA activity of sequences tiling identifies *CTNND2* enhancers. Boxes represent the 25th, 50th, and 75th percentiles. Whiskers extend from hinge to 1.5 times the interquartile range. Data beyond the whiskers are plotted as individual points. Coordinates: hg38. **f-h)** Viral delivery of gRNAs targeting a *CTNND2* TSS and indicated enhancers in HEK293T cells (**f**), as well as two Cri du Chat Syndrome patient cell lines (**g-h**). Bars represent average fold change in *CTNND2* expression (qRT-PCR) relative to control for individual transduction replicates. **i)** Multiplex single-cell CRISPRa screen differential expression test *P*-values for individual *TCF4* TSS- and enhancer-targeting gRNAs. Tracks as in panel **a**. A single isoform of *TCF4* is shown for simplicity. **j)** MPRA activity of sequences tiling identified *CTNND2* enhancers. Coordinates: hg38. Box plots as in panel **d**. **k)** Viral delivery of gRNAs targeting a *TCF4* TSS and indicated enhancer in HEK293T cells. **l)** Viral delivery of gRNAs targeting a humanized *hTcf4* promoter/TSS in *hTcf4+/mTcf4* mouse fibroblasts. Bars represent average fold change in integrated *Tcf4* reporter expression (luciferase). **m-n)** Viral delivery of gRNAs targeting a *TCF4* TSS and indicated enhancer in two Pitt-Hopkins syndrome patient lines). **(k,m-n)** Bars represent average fold change in *TCF4* expression (qRT-PCR) relative to control for individual transduction replicates.

*CTNND2* (5p15.2) encodes the synaptic adhesion protein delta-catenin. Its haploinsufficiency is associated with severe NDDs that phenocopy features of the overlapping Cri-du-chat syndrome (5p deletion)^72–74^. In the multiplex CRIPSRa screen, we identified four TSS-targeting gRNAs and two enhancer-targeting gRNAs that drove upregulation of *CTNND2* in human neurons (**Fig. 5c-e**; **Table S6**). We selected the most effective TSS-targeting gRNA and both enhancer-targeting gRNAs for further validation. Virally mediated targeting of the promoter drove *CTNND2* upregulation in HEK293T cells (3.8-fold, *q* = 0.004), as well as in two iPSC lines derived from individuals with Cri-du-chat syndrome (4.8-fold, *q* = 9e-5; 4.3-fold, *q* = 7e-5; **Fig. 5f-h**). Targeting of the enhancer located 112 kb upstream yielded upregulation of *CTNND2* specifically in the two Cri-du-chat syndrome iPSC lines, but not HEK293T cells (1.8-fold, *q* = 9e-5; 2.1-fold, *q* = 2e-7; **Fig. 5f-h**). Finally, targeting of the enhancer located 2 Mb upstream failed to upregulate *CTNND2* in any of the cell lines, raising the possibility of a false discovery of the multiplex screen, especially in light of its extreme distance (**Fig. 5c, f-h**).

*TCF4* encodes a basic helix-loop-helix transcription factor critical for neurodevelopment^27,75^. Haploinsufficiency of *TCF4* causes Pitt-Hopkins syndrome, a severe NDD associated with ASD, ID, and epilepsy^8,9,75,76^. In the multiplex CRISPRa screen, we identified four gRNAs that drove upregulation of *TCF4*, including three gRNAs targeting alternative TSSs, as well as a gRNA targeting a highly active intronic enhancer (**Fig. 5i-j**; **Table S6**). We selected the most effective TSS-targeting gRNA and both enhancer-targeting gRNAs for further validation. Following viral delivery, we observed modest *TCF4* upregulation with either of these gRNAs in HEK293T cells (1.1-fold, *q* = 0.008; 1.1-fold, *q* = 0.02; **Fig. 5k**). Targeting the *TCF4* promoter in fibroblasts from a transgenic mouse bearing a humanized *TCF4* promoter at the endogenous mouse *Tcf4* locus (*hTcf4*) also yielded upregulation (1.3-fold; *q* = 0.01; **Fig. 5l**; **Fig. S5d-e**). Finally, we observed *TCF4* upregulation in either of two iPSC lines derived from individuals with Pitt-Hopkins syndrome, following viral delivery of either the TSS-targeting (1.6-fold, *q* = 8e-5; 1.5-fold; *q* = 2e-7) or enhancer-targeting (1.2-fold, *q* = 3e-4; 1.1-fold; *q* = 0.02) gRNA (**Fig. 5m-n**).

Overall, in evaluating virally mediated delivery of CRT reagents, we observed the expected upregulation with 7 of 8 (88%) gRNAs (4/4 promoter-targeting and 3/4 enhancer-targeting) in diverse models of *CHD8*, *TCF4*, and *CTNND2* haploinsufficiency, including patient cell lines, human cerebral organoids, and humanized mice (**Fig. 5**; **Fig. S5**; **Table S6**). Taken together, these results demonstrate that many of the gRNAs identified in our CRISPRa screen may serve as effective reagents for vector-mediated upregulation of haploinsufficient NDD risk genes.

## Discussion

Here, we report large-scale functional screens of cCREs for NDD risk genes in human neurons. Our experiments identified thousands of MPRA-active and hundreds of CRISPRa-responsive cCREs that upregulated 200 of 337 prioritized haploinsufficient NDD risk genes, including 91 novel neural enhancer–gene pairs. In follow-up assays, selected cCREs drove tissue-specific activity in the developing mouse CNS and enabled upregulation of well-established NDD risk genes in haploinsufficient patient-derived cell lines, human cerebral organoids, and humanized mice. To our knowledge, this represents the largest compendium of target-linked neural enhancers generated to date, and a foundational resource that can be leveraged toward the development of CRTs at scale.

At this scale and resolution, several consistent patterns of CRISPRa activity in neurons emerged. First, activation was highly specific: targeting of individual cCREs generally upregulated a single gene, rather than broadly affecting all nearby loci, and for distal cCREs, often did so while skipping intervening genes. Second, this specificity extended to promoters, in that most genes exhibited only one or two “activatable” TSSs. Third, even when successful, activation was constrained in magnitude at endogenous promoters and enhancers. Together, these observations indicate that simply targeting CRISPRa near a TSS or within an accessible region is insufficient to achieve robust activation. Instead, successful activation may depend on the capacity of the “pre-targeting” chromatin environment to recruit other factors required for promoter or enhancer activity.

We identified gRNAs and CRISPRa-responsive cCREs supporting upregulation of 200 of 337 (59%) haploinsufficient NDD risk genes. Can these be leveraged for CRT? Therapeutic upregulation for haploinsufficient disorders ideally results in a two-fold increase of the target gene—doubling output from the intact allele to compensate for the loss of the other. Many of our gRNAs approached this mark even without tailoring dose, sequence or activator, *e.g.* 162 of 509 (32%) activating sgRNAs led to a 1.5- to 2.5-fold increase in target gene expression in human neurons. Importantly, even partial restoration might result in therapeutic benefit in some haploinsufficient disorders^21,28^, while overshooting might be buffered by endogenous regulatory feedback. Furthermore, once a responsive CRE has been identified, the precise level of upregulation can be tuned, *e.g.* with gRNA mismatches^77^ or alternative activators^78,79^.

A key consideration is that haploinsufficient genes are, by definition, dose-sensitive—but this sensitivity can cut both ways. Many are also triplosensitive, meaning that excessive expression can itself be deleterious^80,81^. This further underscores the appeal of *cis*-regulatory approaches. As shown here, targeting endogenous promoters or enhancers with CRISPRa yields moderate increases in expression—unlike conventional cDNA-based gene therapy, where heterologous promoters can drive non-specific and often supraphysiologic overexpression. Neural enhancers may be particularly compelling CRT targets, combining cell-type specificity with effect sizes that align with the therapeutic window for dose-sensitive genes^20,22,26^.

Although only a modest proportion of targeted elements yielded upregulation (distal cCREs: 91/2,422 (4%); TSSs: 206/761 (27%)), these hit rates are broadly consistent with previous large-scale noncoding screens—*e.g.* CRISPRi targeting of distal cCREs (2-8% vs. 4% here; **Fig. S6**)^31,82–84^ and CRISPRa targeting of canonical TSS (70% vs. 59% gene-level success here)^23^. Several factors likely contribute to this modest yield. Not all distal cCREs are bona fide enhancers, and not all proximal cCREs correspond to active TSSs. Statistical power also remains a limiting factor, given the cost and sparsity of single-cell profiling; although here we mitigate this 10-fold with multiplexing, false negatives are inevitable. As single-cell technologies continue to scale and decrease in cost, we anticipate higher sensitivity and hit rates in future screens.

We also note key differences that may contribute to a “performance gap” between CRISPRa and CRISPRi. First, activation generally imposes more stringent endogenous requirements (*e.g.* permissive chromatin states, recruitment of cofactors, compatibility with transcriptional machinery, modestly higher target gene baseline expression), whereas repression may be less context-dependent and therefore more uniformly effective^20,22,31,36,85,86^. Second, the extent of activation may be further limited by endogenous feedback mechanisms, in contrast to repression or silencing, which may be less constrained. Finally, gRNA design rules for CRISPRa remain less mature than for CRISPRi^23,56,87,88,86^, though new large-scale datasets (*e.g.* this study, ref ^23^) should accelerate progress toward closing this gap. Together, these factors—combined with the aforementioned points about limited statistical power—suggest that many of the NDD risk genes that were not successfully upregulated (137/337) here may still be targetable, whether through improved gRNA design, more sensitive detection, or the targeting of additional cCREs.

These data also provide a valuable reference point for enhancer–gene prediction models. As described above, the likelihood that a cCREs or enhancer-gene pair would validate by MPRA or CRISPRa was substantially higher when nominated by multiple datasets. ABC scores generated from bulk fetal PFC or iPSC-derived neurons exhibited the strongest positive predictive value: ∼43% of the enhancer-gene pairs validated here were among those predicted by each of these studies^35–37^. However, only 2-3% of all enhancer-gene pairs nominated by these same two studies were experimentally confirmed when tested here. As such, there remains considerable room for improving precision in predicting enhancer-gene pairings. Of note, most existing prediction models have relied on data from a handful of CRISPRi screens performed in transformed cancer cell lines^31,82–84^ for either training or evaluation (**Fig. S6**). By contrast, our data provide functional readouts in differentiated, iPSC-derived post-mitotic neurons, offering a complementary and potentially more relevant foundation for refining enhancer–gene prediction across *in vivo* human cell types.

We conducted our CRISPRa screen using a derivative of *Streptococcus pyogenes* Cas9—the most widely used CRISPR protein—to leverage the extensive ecosystem of associated tools (*e.g.,* inducible CRISPRa machinery, gRNA libraries, and optimized design rules). While SpdCas9 fused to VPH is unlikely to emerge as the ideal configuration for clinical deployment, largely due to its longer length and potential immunogenicity, our results highlight that the most challenging preclinical step of CRT—identifying targetable cCREs—is independent of the effector. Once a functional enhancer or promoter has been mapped, it can be retargeted with alternative Cas variants or activation domains. For example, 34% of our active gRNAs, collectively targeting 109 NDD risk genes, are also compatible with *Staphylococcus aureus* Cas9 PAMs, and in the case of *SCN2A*, nearly identical gRNAs were effective when packaged and delivered with AAV SadCas9-VP64^28^ (**Fig. S5b**). For the remainder, design of new gRNAs for the Cas and activator best suited to therapeutic translation can focus on validated CREs. This separation of target identification and reagent development may facilitate the adaptation of CRT strategies across different effector platforms and delivery systems.

A defining feature of CRT is that it leverages the noncoding genome as a therapeutic substrate. Because so many NDDs arise from haploinsufficiency, this enables the simultaneous discovery of hundreds of targetable elements and matched reagents within a single study. While the translation of each CRT to the clinic remains a major undertaking, the identification of hundreds of validated human neural CREs and gRNAs represents a major step toward that goal and illustrates how at least the groundwork of CRT development can potentially be conducted across many genes and diseases at once. In particular, knowing that upregulation is possible derisks the first steps of a translational pathway, a critical activity in rare genetic disorder therapeutics.

More broadly, only a handful of neural enhancers had previously been functionally mapped to their target genes; here, we substantially expand that set by validating close to one hundred enhancer–gene pairs in human neurons. This resource thus greatly expands the repertoire of experimentally confirmed neural enhancers and provides a foundation for systematic dissection of the regulatory circuits underlying neurodevelopmental disease. Our work necessarily focused on a single neuronal context, and future studies should examine how enhancer responsiveness varies across cell types or subtypes, developmental stages, and disease states. Nonetheless, this study demonstrates the power of multiplex functional genomic screening to chart *cis*-regulatory landscapes in disease-relevant cells. By mapping active enhancers and their linked target genes at scale, we not only advance the prospects for *cis*-regulation therapy but also deepen our understanding of the circuitry that governs human neural gene regulation.

## Supplementary Figures

**Figure S1.**
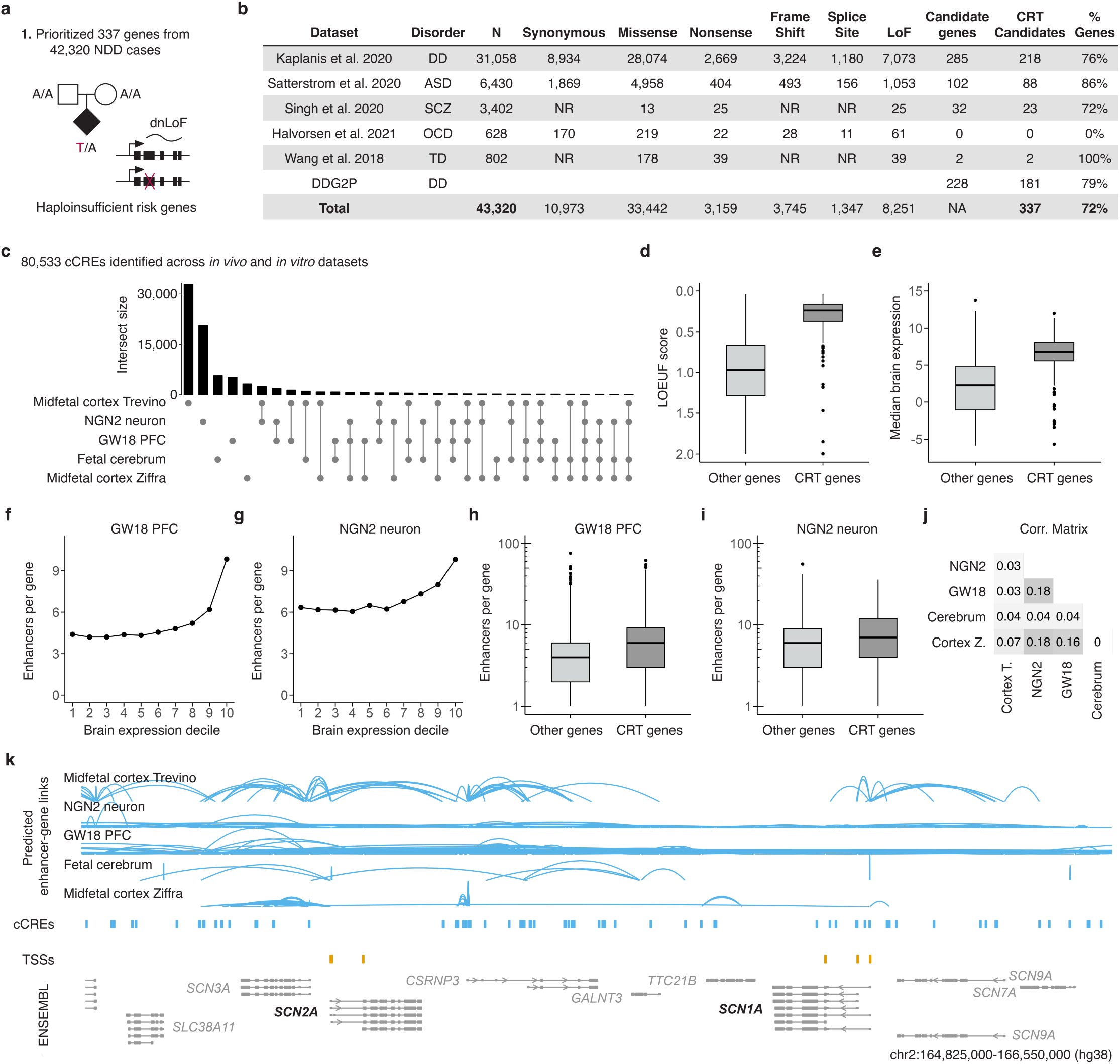
Prioritization of haploinsufficient NDD risk genes and prediction of candidate enhancers. **a-b)** Prioritization of 337 NDD risk genes from 42,320 cases, sources, and variant counts by class. **c)** Loss-of-function observed/expected upper bound fraction (LOEUF) metrics for 337 prioritized NDD risk genes (labeled CRT) vs. all other genes. **d)** Median brain expression for 337 prioritized NDD risk genes vs. all other genes. **e)** Upset plot illustrates the sources of the 80,533 cCREs identified across *in vitro* and *in vivo* data sets. **f)** Predicted enhancers per gene plotted against deciles of predicted target gene expression in gestational week 18 prefrontal cortex (GW18 PFC). **g)** Predicted enhancers per gene plotted against deciles of predicted target gene expression in iPSC-derived NGN2 neurons. **h)** Predicted enhancers per gene for 337 prioritized NDD risk genes vs. all other genes in GW18 PFC. **i)** Predicted enhancers per gene for 337 prioritized NDD risk genes vs. all other genes in iPSC-derived NGN2 neurons. **j)** Spearman correlation across enhancer prediction approaches. Correlations < 0.005 round to 0 in the visualization. **k)** Tracks for predicted cCREs (blue) and linked TSSs (orange) for all genes in the indicated genomic range are visualized alongside tracks for RefSeq validated transcripts (ENSEMBL/NCBI). Prioritized NDD risk genes are labeled in black.

**Figure S2.**
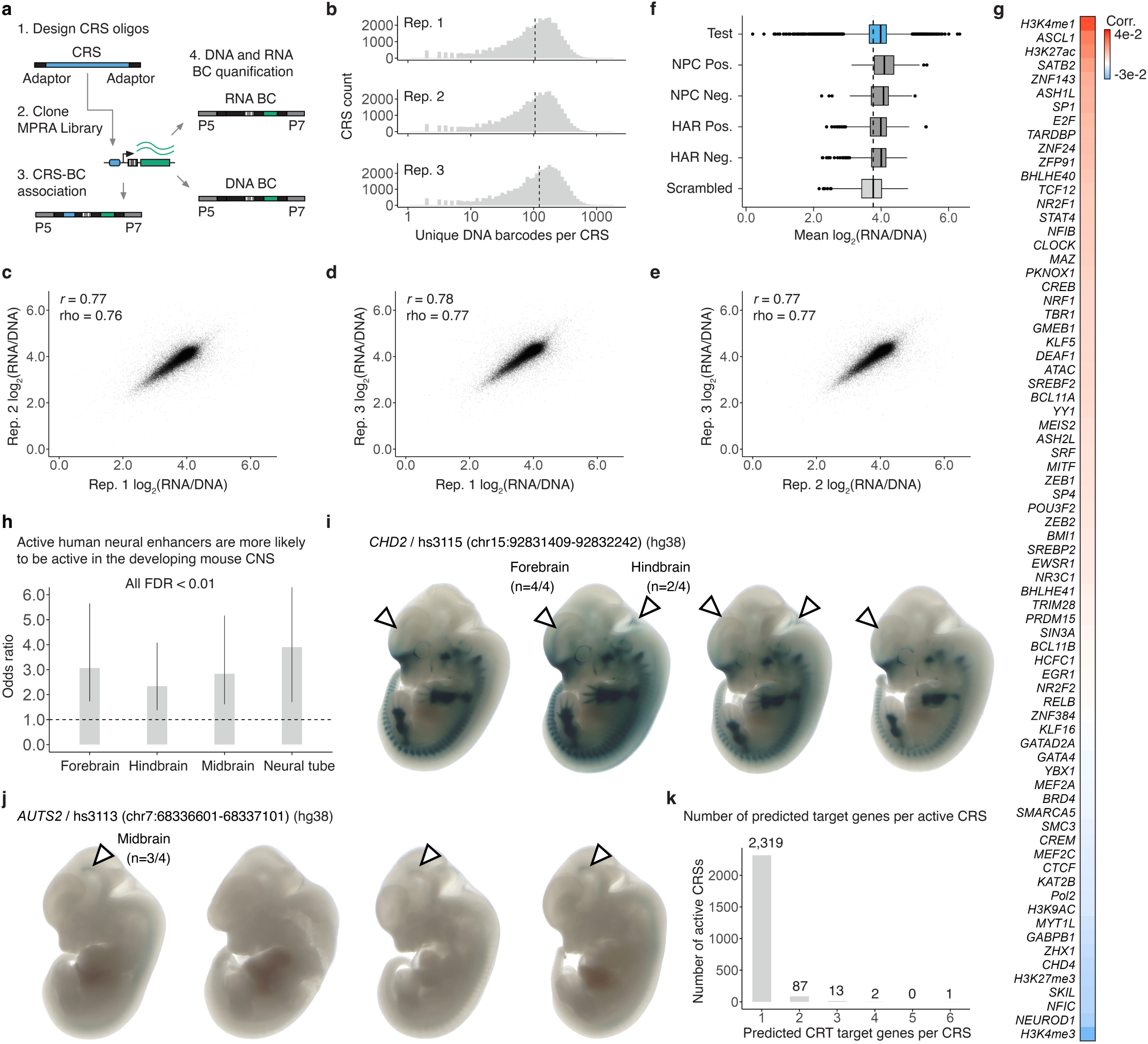
MPRA design, quality control, and additional results. **a)** MPRA cloning and sequencing workflow schematic. **b)** Number of unique DNA barcodes recovered per CRS across three transduction replicates. Dashed line represents the median. **c-e)** MPRA activity score correlations across transduction replicates. **f)** MPRA activity score of CRSs grouped by source (**Methods**). Dashed line represents the median of scrambled control sequences. **g)** Correlation between MPRA activity scores and 74 epigenetic/chromatin feature scores derived from neural samples. **h)** Odds ratio of developing mouse CNS reporter expression given MPRA activity. **i-j)** Reporter expression of two MPRA active human enhancers in the developing mouse CNS. Fractions indicate proportion of profiled embryos with detectable expression in the indicated CNS tissue. **k)** Number of predicted target genes per active CRS.

**Figure S3.**
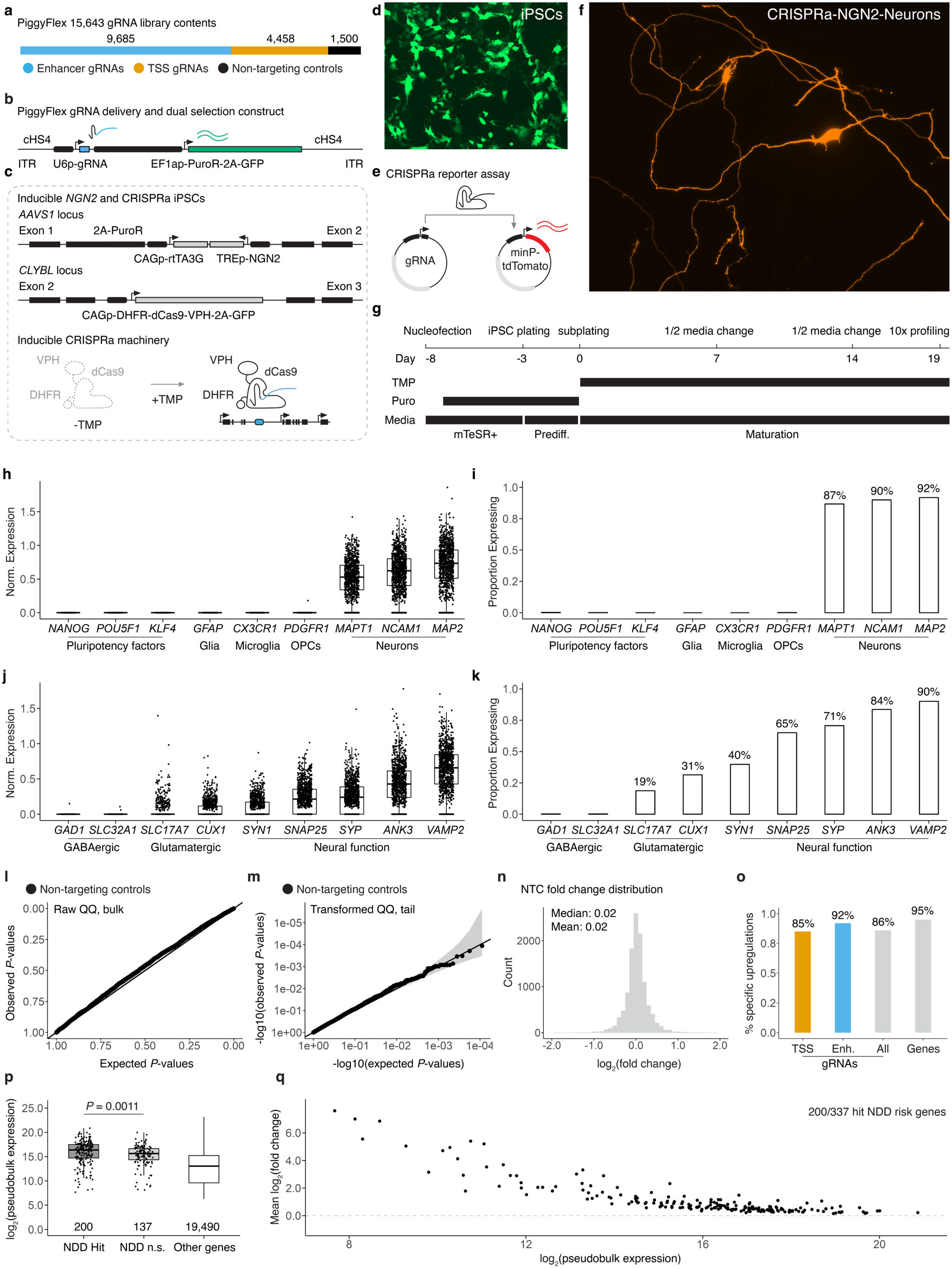
Multiplex single cell CRIPSRa screen design, quality control, and additional results. **a)** PiggyFlex gRNA library contents. **b)** PiggyFlex gRNA delivery and dual selection construct. **c)** Integrated, inducible *NGN2* and CRISPRa transgene schematics. Dox-inducible *NGN2* drives neural differentiation from iPSCs. TMP stabilizes/induces functional CRISPRa machinery. **d)** Selected iPSCs expressing piggyFlex gRNA construct GFP. **e)** minP-tdTomato CRISPRa reporter assay schematic. When co-transfected with the gRNA and reporter plasmids, cells with functional CRISPRa machinery drive strong expression of the otherwise lowly expressed minP-tdTomato reporter. **f)** Differentiated CRISPRa-*NGN2* iPSC-derived neurons transfected with the minP-targeting gRNA and minP-tdTomato reporter drive strong tdTomato expression. **g)** Screen differentiation, selection, and profiling timeline. **h)** Expression of marker genes. Dots represent normalized expression from individual single-cell transcriptomes. Cells are downsampled to 1000 cells for visualization. **i)** Proportion of cells with detectable expression of marker genes. Proportions are calculated on the full set of single cell transcriptomes. **j)** Expression of marker genes. Visualization as in panel **h**. **k)** Proportion of cells with detectable expression of marker genes. Visualization as in panel **i**. **l-m)** Raw and transformed quantile-quantile plots showing distribution of expected vs. observed *P*-values for NTC differential expression tests. **n)** NTC differential expression fold change distribution. **o)** Proportion of upregulations that were target gene specific - i.e. yielded upregulation of only one gene within 1Mb of the target site in *cis*. The first three bars indicate the proportion of TSS-targeting gRNAs (orange), enhancer-targeting gRNAs (blue) and all gRNAs (gray) yielding specific upregulations. The last bar indicates the proportion of the 200 activatable NDD risk genes for which there were one or more specific activating gRNAs identified (190/200, 95%). **p)** Pseudobulk expression levels of NDD risk genes for which an activating gRNA was identified (NDD hit) or not (NDD n.s.). *P*-value is from a Wilcoxon rank-sum test. NDD risk genes are plotted next to all other expressed genes. **q)** Expression level (x-axis) plotted against the average log2 fold-change across activating gRNAs for the 200/337 NDD risk genes.

**Figure S4.**
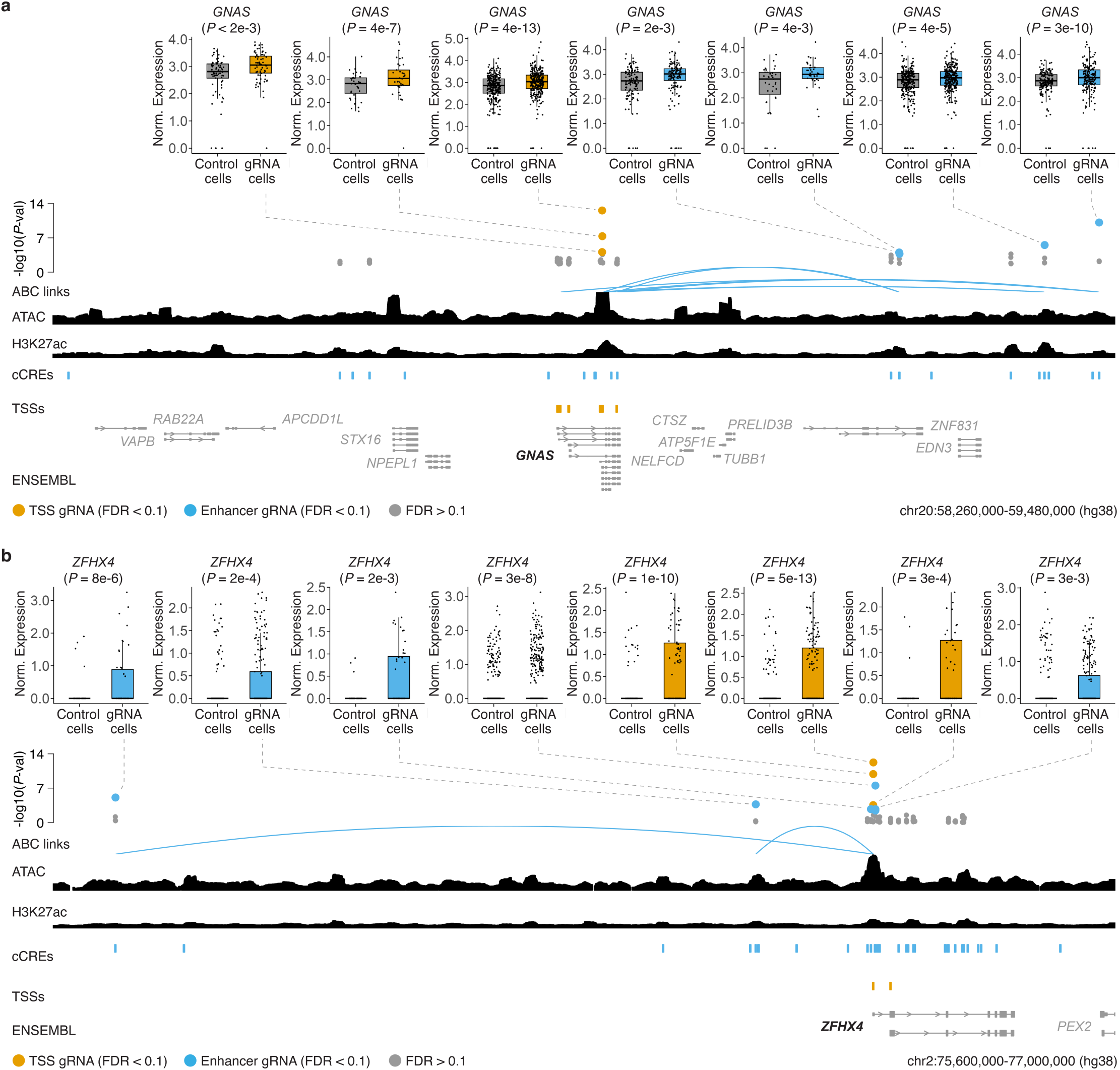
Highlighted examples of MPRA-active and CRISPRa-responsive enhancers at NDD risk loci. **a)** (top) Box plots showing log2 fold-change in *GNAS* expression for gRNA cells compared to control cells. Dots represent normalized expression from individual single cell transcriptomes. Control cells are downsampled to have the same number of cells as the indicated targeting gRNA for visualization. (bottom) Multiplex single cell CRISPRa screen differential expression test *P*-values for individual *GNAS* TSS- and enhancer-targeting gRNAs are plotted above tracks for predicted cCREs (blue) and linked TSSs (orange) for prioritized NDD risk genes. TSS- and enhancer-targeting gRNAs yielding significant target gene upregulation (FDR < 0.1) are coloured orange or blue, respectively. These results are plotted above tracks for iPSC-derived neuron ATAC-seq^35^, H3K27ac^35^, and RefSeq validated transcripts (ENSEMBL/NCBI). Here, only predicted enhancer gene links that were supported by both MPRA and multiplex single cell CRISPRa screens are shown. **b)** *ZFHX4* locus results. Tracks as in panel **a**.

**Figure S5.**
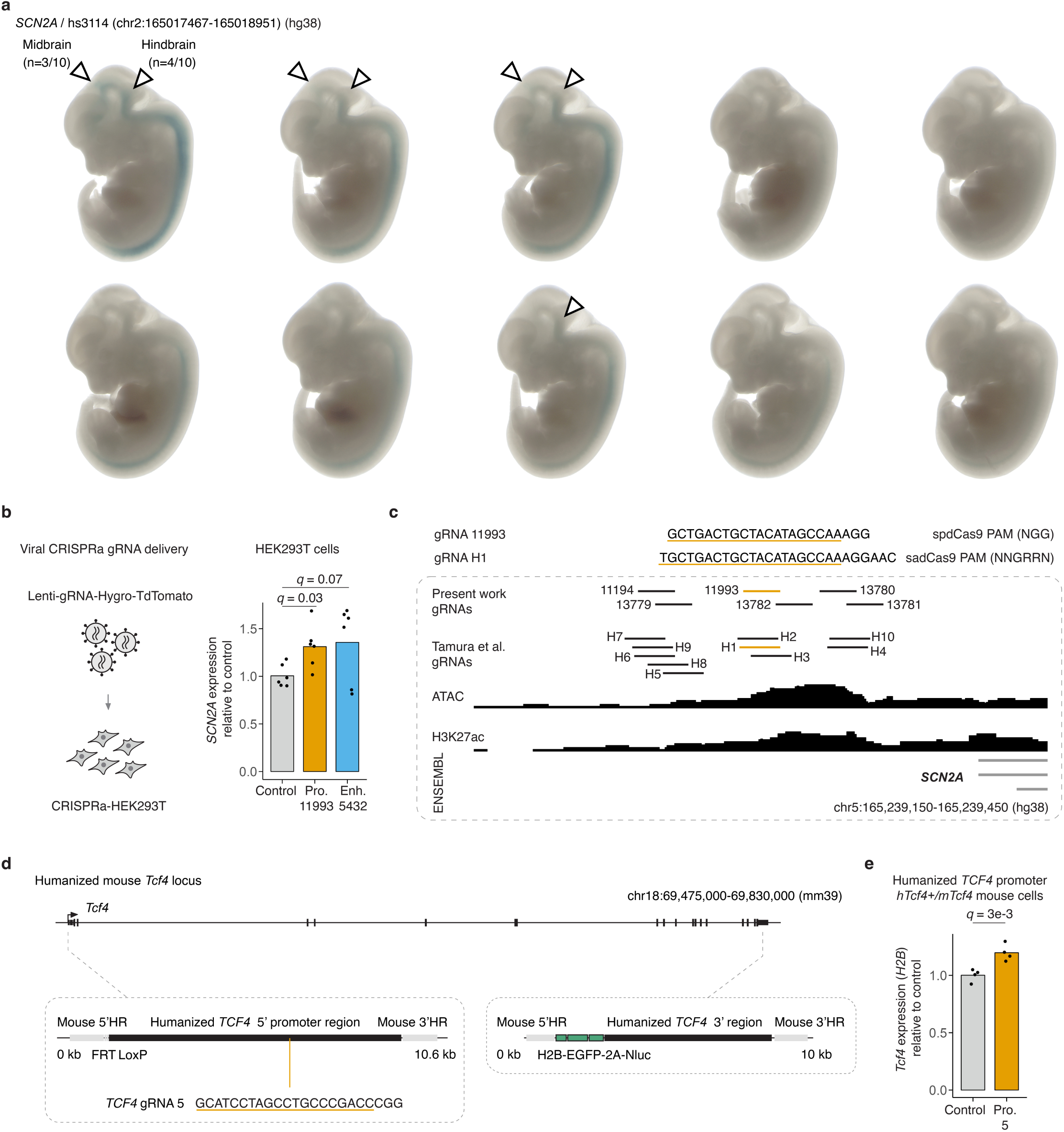
Additional *SCN2A* and *TCF4* activation and reporter experiments. **a)** Viral delivery of gRNAs targeting a *SCN2A* TSS and enhancer in HEK293T cells. Bars represent average fold change in *SCN2A* expression (qRT-PCR) relative to control for individual transduction replicates. **b)** Overlap of *SCN2A* TSS-targeting gRNAs tested in the present large-scale screen and a previous one-by-one study using sadCas9. The two strongest activating gRNAs overlap. **c)** Reporter expression driven by a human *SCN2A* enhancer in the developing mouse CNS. Fractions indicate proportion of profiled embryos with detectable expression in the indicated CNS tissue. **d)** Schematic of the engineered mouse *Tcf4* locus with a humanized *Tcf4* 5’ and 3’ regions. **e)** Viral delivery of a gRNA targeting the *TCF4* TSS in *hTcf4+/mTcf4* mouse fibroblasts with a humanized *TCF4* promoter. Bars represent average fold change in *Tcf4* reporter (*H2B*) expression (qRT-PCR) relative to control for individual transduction replicates.

**Figure S6.**
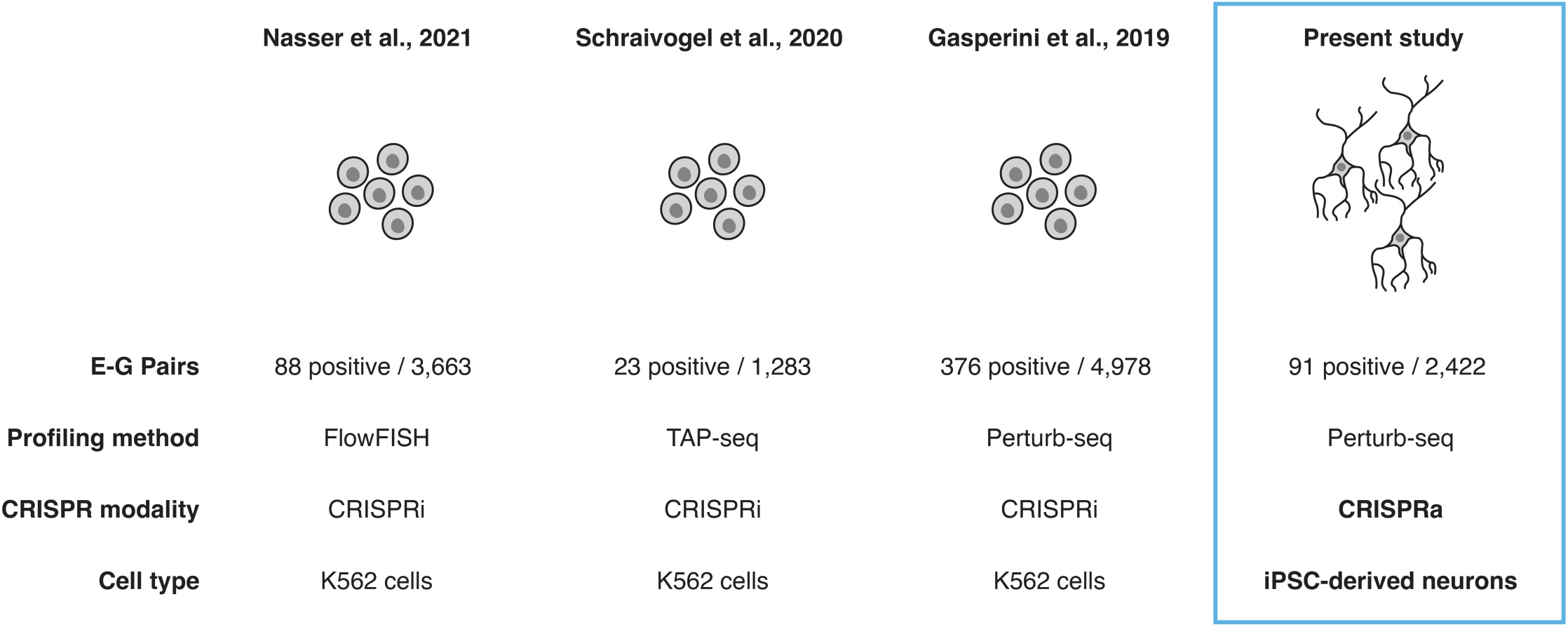
Comparison of large-scale single-cell noncoding CRISPR screens. Comparison of E-G pairs identified out of the number tested for recent large-scale noncoding CRISPR screens. Numbers for the first three CRISPRi datasets collected in K562 cells^31,82,84^ are from a recent reanalysis (ENCODE)^83^. Profiling methods, CRISPR modality, and target cell types are also listed.

## Acknowledgements

We are grateful to numerous members across all collaborating labs for comments, suggestions, and discussions on this work. We thank the technical team at the Lawrence Berkeley National Laboratory for generating transgenic mouse embryos. The Human WTC11 NGN2 ecDHFR-dCas9-VPH line was a kind gift from the M. Kampmann lab at UCSF.

## Funding

This work was supported by the Weill Neurohub (to S.J.S., N.A., and J.S.), the National Human Genome Research Institute (UM1HG011966 to N.A. and J.S.), the National Institute of Mental Health (U01MH122681 and R01MH116999 to S.J.S. and R01MH125246 to N.A.), the National Institute on Aging (R01AG050986 to N.A.) and the Medical Research Council Centre of Research Excellence in Therapeutic Genomics (MR/Z504725/1 to S.J.S and N.A.). M.K and L.A.P were supported by the National Human Genome Research Institute (R01HG003988 to L.A.P.) and conducted at the E.O. Lawrence Berkeley National Laboratory under the US Department of Energy Contract DE-AC02-05CH11231, University of California. T.A.M was supported by a Banting Postdoctoral Fellowship from the Natural Sciences and Engineering Research Council of Canada (NSERC). N.F.P. was supported by a National Science Foundation (NSF) and Autism Science Foundation (ASF) graduate research fellowship. D.C. was supported by award no. F32HG011817 from the National Human Genome Research Institute. D.H.G. was supported by the National Institute of Mental Health (NIMH R01MH10027, U01MH116489, RM1MH132651) and an Eagles Autism Foundation pilot grant. G.T.C was supported by the National Institute of Mental Health (NIMH F32MH124337). J.-B.L. was supported by the Damon Runyon Foundation (DRG-2435-21) and by a Next Generation Scientist award from the Cancer Research Society of Canada (grant no. 1155581). A.R.M. was supported by the National Institutes of Health R01 grant (R01MH123828). B.D.P. was supported by the National Institutes of Health (R01NS129914, R01NS131615, R01NS114086). M.E.T. was supported by the National Institutes of Health (R01HD105266, R01MH115957, R01MH129722, U01HG011755) and the Simons Foundation (SFARI 1009802). J.S. is an Investigator of the Howard Hughes Medical Institute.

## Author contributions

Conceptualization, J.S., S.J.S., and N.A.; Investigation, T.A.M., N.F.P., F.M.C., G.T.C., M.K., L.M.J., and H.C.N.; Data Curation, T.A.M., N.F.P., F.M.C., R.M.D.; Formal Analysis, T.A.M., N.F.P., and F.M.C.; Visualization, T.A.M., N.F.P., and F.M.C.; Resources, N.A. and J.S.; Supervision, S.J.S., N.A. and J.S.; Writing – Original Draft, T.A.M., N.F.P., F.M.C., N.A. and J.S.; Writing – Review & Editing, T.A.M., N.F.P., F.M.C., R.M.D., G.T.C., M.K., L.M.J., H.C.N., D.C.L., A.S.L., P.V., D.C., J.B.L., B.K.M., K.F., M.E.T., A.M., B.P., L.A.P., D.H.G., S.J.S., N.A., and J.S.; Funding Acquisition, S.J.S., N.A., and J.S.

## Competing interests

S.J.S. receives research funding from BioMarin Pharmaceutical Incorporated. N.A. is the cofounder and on the scientific advisory board of Regel Therapeutics and received funding from BioMarin Pharmaceutical Incorporated. J.S. is on the scientific advisory board, a consultant, and/or a co-founder of Prime Medicine, Guardant Health, Camp4 Therapeutics, Phase Genomics, Adaptive Biotechnologies, Sixth Street Capital, Pacific Biosciences, Somite AI and 10x Genomics. All other authors declare no competing interests. Dr. Muotri is the co-founder of and has an equity interest in TISMOO, a company dedicated to genetic analysis and human brain organogenesis, focusing on therapeutic applications customized to autism spectrum disorders and other neurological diseases. The terms of this arrangement have been reviewed and approved by the University of California San Diego, in accordance with its conflict-of-interest policies. UCSD has filed a patent application (WO2022072709A1), in which A.R.M. is an inventor.

## Disclosures

We disclose that language editing and proofreading were supported by AI-based tools; these were not used for conceptual development or primary manuscript writing. The authors take full responsibility for the contents of this manuscript.

## Data Availability

Raw sequencing data, custom sequencing amplicons, and processed data files generated in this study have been deposited on the IGVF portal and are freely available with accession numbers IGVFDS1120GRVG (MPRA), and IGVFDS2290SSEF (single cell CRISPRa screen).

## Code availability

Analysis and visualization code are available at GitHub (https://github.com/shendurelab/Neural_CRE_MPRA_CRISPRa/).

## Methods

### Risk gene prioritization

We began by compiling a list of 466 genes implicated in NDD. This includes 285 genes associated with DD at genome-wide significance^9^, 102 genes associated with ASD at FDR≤0.1^8^, 32 genes associated with schizophrenia at FDR≤0.1^16^, two genes associated with Tourette disorder at FDR≤0.1^18^, and 228 clinician-curated genes implicated in monoallelic loss-of-function (LoF) brain disorders from the Decipher Developmental Disorder Genotype – Phenotype Database (DDG2P)^93^ (**Fig. S1**). To distinguish genes with LoF mechanisms that are ideal candidates for CRT, we compiled *de novo* variants from 42,320 offspring with parental data to exclude neurodevelopmental or neuropsychiatric risk genes with fewer observed variants than expected based on sample size, gene length, and sequence composition. This includes 31,058 developmental disorder^9^, 6,430 autism spectrum disorder^8^, 3,402 schizophrenia^16^, 802 Tourette disorder^18^, and 628 obsessive compulsive disorder^17^ offspring. A total of 8,251 *de novo* protein truncating variants, including 3,745 frameshift, 1,347 canonical splice site, and 3,159 stop gain variants, were compared to expectation, in addition to 33,442 missense and 10,973 synonymous variants as calculated by the denovolyzeR tool^32,94^. As expected, there was an enrichment of observed/expected (O/E) *de nov*o missense (O/E=1.22, p<0.001) and LoF variants (O/E=2.17, p <0.001) in our neurodevelopmental or neuropsychiatric disorder cohorts. 337 genes had a ratio of O/E *de novo* LoF variants >1 and were prioritized as candidates for CRT.

### cCRE prediction and prioritization

We combined enhancer-gene predictions from five datasets characterizing the regulatory landscape of the developing human brain, together with various mapping strategies (**Fig. 1c**; **Fig. S1e**; **Methods**). These were: 1) Paired sc-RNA-seq and sc-ATAC-seq data from human midfetal cortex (gestational week (GW) 18-26) and correlation-based mapping^33,34^; 2) Bulk ATAC-seq data from iPSC-derived excitatory neurons (differentiated for 7-8 weeks; *NGN2* neurons, as used later in this study) and Activity-By-Contact (ABC) mapping^35,36^; 3) Bulk ATAC-seq data from human midfetal prefrontal cortex (GW18) and ABC mapping^36,37^; 4) Single cell ATAC-seq data from human midfetal cerebral excitatory neurons (GW14-20) and Cicero co-accessibility mapping^38,39^; and 5) Single cell ATAC-seq data from human midfetal cortex excitatory neurons (GW17-21) and ABC mapping^36,40^. Enhancer-gene predictions from GW18 PFC and *NGN2* neurons were converted to hg38 using UCSC liftOver and each dataset was filtered for pairs that included one of our 337 CRT candidate genes. All cCREs were extended from their center to be at least 500bp wide and then combined using bedtools merge. If a cCRE overlapped with a 2000bp promoter region or an exon of a protein coding transcript it was removed. After merging and filtering there were a total of 5,425 unique cCREs.

### Construct design and library cloning

#### MPRA enhancer tiling library

We designed a MPRA library consisting of 44,312 x 270 bp sequences densely tiling all 5,425 cCREs with 90 bp overlaps. We also included 100 active and 99 inactive control sequences from an MPRA of open chromatin regions conducted in neural precursor cells (NPCs)^47^, 729 active and 600 inactive control sequences from an MPRA of human accelerated regions (HARs) conducted in an SH-SY5Y immortalized neuroblastoma cell line^48^, and base-level shuffled versions of a randomly selected subset of the 44,312 tiles. Altogether, the lentiMPRA library included 46,340 unique designs (**Table S3**).

The MPRA library was synthesized by Twist Bioscience as an oligo pool. Pools were resuspended in IDTE buffer pH 8.0 (IDT, Cat. No. 11-05-01-13) and amplified with custom 5BC-AG-f01 and 5BC-AG-r01 primers using Kapa HiFi Hotstart readymix (KAPA Biosystems, Cat. No. 08202940001; cycling conditions: 98°C for 3 minutes, 5 cycles of 98°C x 20 seconds, 65°C x 15 seconds, and 72°C x 15 seconds, and 72°C x 1 minute). The PCR product was combined in a DNA low bind tube and purified with 0.85x SPRI select beads (Beckman Coulter, Cat. No. B23318). DNA was eluted in 50 uL EB and then a 2nd PCR reaction was performed with 5BC-AG-f02 and 5BC-AG-r02 primers to add 15 bp transcribed MPRA barcodes using Kapa HiFi Hotstart readymix (KAPA Biosystems, Cat. No. 08202940001; cycling conditions: 98°C for 3 minutes, 10 cycles of 98°C x 20 seconds, 65°C x 15 seconds, and 72°C x 15 seconds, and 72°C x 1 minute). The PCR product was combined in a DNA low bind tube and purified using 0.75x SPRI beads.

pLS-SceI backbone vector (Addgene, Cat. No. 137725) was linearized by restriction enzyme digestion with AgeI-HF (NEB, Cat. No. R3552S) and SbfI-HF (NEB, Cat. No. R3642S) in CutSmart buffer for 1 hour at 37°C. Digest was purified with 0.65x SPRI beads and eluted in 50 uL EB. Amplified oligos from PCR2 were recombined with linearized pLS-SceI backbone via gibson assembly. Briefly, 208 fmol of linearized pLS-SceI was recombined with 965 fmol of purified DNA insert using NEBuilder Hifi DNA assembly master mix (NEB, Cat. No. E2621S). Recombination product was purified using 0.65x SPRI beads and eluted with 45 uL EB. To remove the uncombined background fragments, an additional restriction enzyme digest with I-SceI (NEB, Cat. No. R0694S) in CutSmart buffer for 1 hour at 37°C.

High complexity MPRA libraries were electroporated in NEB 10-beta competent *E. coli* (NEB, Cat. No. C3019H) using a Gemini X2 electroporator (BTX, Cat. No. 452006) and the C3020 program (parameters: voltage, 2kV; resistance, 200 ohms; capacitance, 2 uF; number of pulses, 1; gap width, 1 mm). Prewarmed SOM media was immediately added following electroportation and bacteria were transferred to a round bottom culture tube and incubated at 37°C shaking for 1 hour recovery. Transformed bacteria were cultured on pre-warmed 15 cm LB agar plates (Millipore Sigma, Cat. No. L5667) with 400 uL undiluted bacteria and 100 uL of 100 mg/mL carbenicillin per agar plate. Bacteria were cultured overnight in a 37°C incubator and the following day 16 colonies were picked and sent of whole plasmid sequencing (Plasmidsaurus) to confirm library integrity. Bacteria were harvested using a cell scraper and 10 mL of LB and centrifuged at 3,400 x g for 10 min before harvesting plasmid DNA with ZymoPURE II Plasmid Midiprep Kit (Zymo Research, Cat. No. D4201). Association sequencing was performed following plasmid harvest to map cCREs to their corresponding 15 bp transcribed barcodes; we achieved a final library complexity of 9,338,288 barcodes across 44,729 cCREs.

#### piggyFlex CRISPRa sgRNA library

CRISPRa gRNAs targeting the 2,422 enhancers and 761 promoters/TSSs for the 337 NDD risk genes were designed using Flashfry^95^. 1,500 NTC gRNAs and additional TSS-targeting gRNAs were further selected from the Horlbeck et al. CRISPRa-V2 library^56^. In total, the library contained 15,643 gRNAs, including 9,685 enhancer-targeting gRNAs, 4,458 TSS-targeting gRNAs, and 1,500 NTCs (**Fig. S3**; **Table S4**).

Custom R scripts and Flashfry commands were used to design the gRNA library and append sequences for cloning. In brief, NGG protospacers were identified and scored with FlashFry using default parameters^95^. For promoter-targeting gRNAs, a TSS-distance metric was calculated based on human fetal brain 5′ CAGE data from FANTOM (https://fantom.gsc.riken.jp/5/sstar/FF:10085-102B4; CTSS, hg38)^41^. Genome-wide CRISPRa design rules were used to estimate optimal TSS-distances^87^. For enhancers, gRNAs closer to the center of the candidate enhancer sequences were prioritized^86^. gRNAs containing problematic features such as polythymidine stretches or extreme GC content were excluded.

We then performed six iterative rounds of gRNA selection with successively relaxing score and distance thresholds to select the top 4 gRNAs for each TSS and candidate enhancer. Finally, to facilitate Pol III transcription, we modified all gRNAs to begin with a G followed by the 19 base pair spacer sequence. All oligo sequences, including gRNA spacers and appended cloning adapters are listed in **Table S4**.

gRNAs were ordered as an oligo pool from Twist Bioscience with flanking sequences for cloning into piggyFlex. Oligos were double stranded across multiple low cycle PCR reactions using Q5 polymerase (NEB, Cat. No. M0492L; cycling conditions: 98°C for 30 seconds, 5 cycles of 98°C × 10 seconds, 65°C x 15 seconds and 72°C x 30 seconds). PCR products were then pooled and purified using 2.0x AMPure XP beads (Beckman Coulter, Cat. No. A63880), then cloned into piggyFlex via Gibson Assembly using NEBuilder HiFi assembly master mix (NEB, Cat. No. E5520S). Cloned libraries were then electroporated in triplicate into NEB 10-beta electrocompetent E. coli (NEB, Cat. No. C3020K), cultured at 30°C overnight and prepared using a Zymo Pure II (Cat. No. D4200) kit following manufacturer protocols. The resulting library was then amplicon sequenced to confirm gRNA presence and library balance. In brief, gRNA sequencing libraries were generated using a two step PCR process to amplify gRNAs then append sequencing adapters and sample indices. gRNAs were amplified across eight reactions using Q5 polymerase (NEB, Cat. No. M0492L; cycling conditions: 98°C for 30 seconds, 15 cycles of 98°C × 10 seconds, 65°C x 15 seconds and 72°C x 40 seconds). SYBR Green (Thermo Fisher Scientific, Cat. No. S7567) was added to track the amplification curve. PCR products were pooled and purified using 1.1x AMPure XP beads (Beckman Coulter, Cat. No. A63880). Sequence flow cell adapters and dual sample indices were then appended in the second PCR reaction using Q5 polymerase (NEB, Cat. No. M0492L; cycling conditions: 98°C for 30 seconds, 5 cycles of 98°C × 10 seconds, 65°C x 15 seconds and 72°C x 30 seconds). PCR products were purified using 0.9x AMPure XP beads (Beckman Coulter, Cat. No. A63880) and assessed on an Agilent 4200 TapeStation before sequencing on an Illumina NextSeq 2000. Following sequencing confirmation, the library was double digested with SapI (NEB, Cat. No. R0569L) and the piggyFlex^22^ puromycin and GFP selection cassettes were inserted via Gibson assembly with NEBuilder HiFi assembly master mix (NEB, Cat. No. E5520S). The resulting final piggyFlex library was transformed in quadruplicate to NEB 10-beta electrocompetent *E. coli* (NEB, Cat. No. C3020K), prepared using a Zymo Pure II kit (Cat. No. D4200), then pooled and amplicon sequenced (as above) to reconfirm gRNA presence and balance.

Primer sequences are provided in **Table S7**. gRNA sequences are provided in **Table S4**.

#### Lentiviral gRNA constructs

Single stranded DNA oligos were ordered from IDT with forward and reverse complement sgRNA sequences FWD: 5’-CACCG(*sgRNA_sequence*)-3’, REV: 5’-AAAC(*reverse_compliment_sgRNA_sequence*)C-3’. Ligation reaction was performed using T4 PNK (NEB, Cat. No. M0201), 10 mM ATP (Thermo Fisher Scientific, Cat. No. 2706649), and forward and reverse sgRNA oligos (Cycling conditions: 37°C x 30 minutes, 95°C x 5 minutes, and then cooled with a ramp speed of 5°C/minute to 25°C). Annealed sgRNA oligos were diluted 1:10 and cloned into the *lentiGuide-Hygro-dTomato* (Addgene, Cat. No. 99376) or *pLV hU6-gRNA(anti-sense) hUbC-VP64-dCas9-VP64-T2A-GFP* (Addgene, Cat. No. 66707) lentiviral backbone via golden gate reaction with BsmBI-V2 (NEB, Cat. No. R0739S) and T4 ligase (NEB, Cat. No. B0202S) (Cycling conditions: 15x cycles of 37°C for 5 minutes, 23°C for 5 minutes, followed by hold at 4°C). Competent *E. coli* (NEB, Cat. No. C3020K) were transformed using 1 uL of golden gate product and plated on ampicillin resistance agar plates (Millipore Sigma, Cat. No. L5667). Single colonies were picked the following day to inoculate minicultures and plasmid DNA was harvested using QIAprep Spin Miniprep Kit (Qiagen, Cat. No. 27104). Cloned gRNA constructs were confirmed via whole plasmid sequencing (Plasmidsaurus) prior to lentivirus production. Targeting sgRNAs for *CTNND2* and *TCF4* were compared to a scrambled sgRNA with the following sequence: *GGCTCCTGCCACAGTCCCCA*.

### Cell lines and culture

#### HEK293T and lentivirus production

Lenti-X HEK293T (Takara, Cat. No. 632180) cells were cultured in DMEM (ThermoFisher Scientific, Cat. No. 11995-065) supplemented with 10% heat-inactivated FBS (ThermoFisher Scientific, Cat. No. A5669501) and 1% PenStrep (ThermoFisher Scientific, Cat. No. 15140-122) and underwent media changes every 2-3 days. After reaching 80-90% confluency, cells were routinely passaged following dissociation with TrypLE Express (ThermoFisher Scientific, Cat. No. 12604-013). For lentivirus production, ∼3 million cells were plated on a 15 cm dish 2 days prior to transfections. Cells were transfected with 6.4 ug lentiviral construct and 12.8 uL HIV packaging mix using 38.4 uL EndoFectin transfection reagent (Genecopoeia, Cat No. LT002). ∼12-16 hours following transfection, a media change was performed to DMEM (ThermoFisher Scientific, Cat. No. 11995-065) supplemented with 2% heat-inactivated FBS (ThermoFisher Scientific, Cat. No. A5669501), 1% PenStrep (ThermoFisher Scientific, Cat. No. 15140-122), and TiterBoost reagent (Genecopoeia, Cat No. LT002). Approximately 48 hours post transfection, media supernatant containing lentiviral particles was harvested in 50 mL falcon tubes and cellular debris was pellet by spinning at 1000 x g for 10 minutes at 4°C. Remaining media was passed through a 0.45 um PES filter (ThermoFisher Scientific, Cat. No. 166-0045) and ⅓ volume lenti-X concentrator (Takara, Cat. No. 631231) was added to filtered supernatant and allowed to cool for at least 48 hours to precipitate lentiviral particles. Harvesting was performed by centrifugation at 1500 x g for 45 minutes at 4°C followed by discarding supernatant and an additional spin at 1500 x g for 5 minutes at 4°C. Supernatant was removed once more and lentivirus-containing pellets were resuspended in 300 uL 1x PBS per 15 cm plate used for lentivirus production. Harvested virus was frozen at -80°C in 50 uL aliquots until use. For CRISPRa validation experiments, lentivirus was titered via qRT-PCR prior to use (Genecopoeia, Cat No. LT005).

To generate stable CRISPRa expressing HEK293T line, cells were transduced at high MOI with *lenti-EF1a-dCas9-VP64-Puro* (Addgene, Cat. No. 99371) and incubated for 3 days to allow for lentiviral integration and expression. Following lentivirus expression, cells underwent 1ug/mL puromycin selection for 3 days and were allowed to recover for at least 3 days. Strong Sp-dCas9 expression relative to untransduced HEK293T was confirmed via qRT-PCR prior to starting experiments.

#### iPSC-derived neurons for MPRA

NGN2-iNeurons were derived from WTC11 iPSCs containing a doxycycline-inducible *NGN2* transgene integrated into the *AAVS1* locus^49^. Cells were obtained from the laboratory of Dr. Yin Shen at the University of California, San Francisco (UCSF). iPSCs were maintained in StemFlex medium (ThermoFisher Scientific, A3349401) and underwent daily media changes. Cells were routinely passaged following dissociation with accutase (ThermoFisher Scientific, A11105-01) and replated on matrigel (Corning, 356231) coated plates. Media was supplemented with 10 µM ROCK inhibitor Y-27632 (Tocris, 1254) on the day of plating. For pre-differentiation, cells were replated on matrigel coated plates and cultured for 3 days in KnockOut DMEM/F−12 (ThermoFisher Scientific, 12660-012) medium supplemented with 2 µg/mL doxycycline (Sigma-Aldrich, D3447-500MG), 1x N-2 Supplement (ThermoFisher Scientific, 17502-048), 1x NEAA (ThermoFisher Scientific, 11140-050), 10 ng/mL NT-3 (PeproTech, 450-03), 10 ng/mL BDNF (PeproTech, 450-02), and 1 µg/mL mouse laminin (ThermoFisher Scientific, 23017-015). Cells underwent daily media changes during pre-differentiation and 10 µM ROCK inhibitor Y-27632 was supplemented on the first day of plating. Following 3 days of pre-differentiation, cells were dissociated with accutase and plated on Poly-L-Ornithine (Sigma-Aldrich, P3655-10MG) coated 10 cm plates. NGN2-iNeurons were cultured in BrainPhys neuronal media (StemCell Technologies, 5790) supplemented with 2 µg/mL doxycycline, 1X N-2 Supplement, 0.5X B-27 Supplement (ThermoFisher Scientific, 12587-010), 1X NEAA, 10 ng/mL NT-3, 10 ng/mL BDNF, and 1 µg/mL mouse laminin. A half-media change was conducted on day 7 without doxycycline.

#### iPSC-derived neurons for multiplex, single cell CRISPRa screening

WTC11 iPSCs harboring a doxycycline-inducible *NGN2* transgene at the *AAVS1* safe-harbor locus^49^ and an ecDHFR-dCas9-VPH construct (VPH = 12×VP16 fused to P65-HSF1)^96^ at the *CLYBL* safe-harbor locus were a gift from Martin Kampmann^58^ (Fig. S3). Cells were maintained in mTeSR+ Basal Medium (Stemcell Technologies, Cat. No. 100-0276) on Greiner Cellstar plates (Sigma-Aldrich) pre-coated with Geltrex™ LDEV-Free hESC-Qualified Basement Membrane Matrix (Gibco, Cat. No. A1413302) diluted 1:100 in Knockout DMEM (Thermo Fisher Scientific, Cat. No. 10829018). Medium was refreshed every other day. At ∼70–80% confluence, cells were passaged by aspirating medium, rinsing with DPBS (Thermo Fisher, Cat. No. 14190144), treating with Accutase (Thermo Fisher, Cat. No. A1110501) for 5 min at 37 °C, quenching 1:1 with mTeSR+, centrifuging at 500 g for 3 min, and resuspending in mTeSR+ with 0.1% ROCK inhibitor (Stemcell Technologies, Cat. No. Y-27632). Cells were counted and replated on Geltrex-coated plates at the desired density.

For neuronal differentiation, selected iPSCs harboring piggyFlex gRNAs were dissociated using accutase and resuspended in predifferentiation medium consisting of Knockout DMEM/F-12 (Thermo Fisher, Cat. No. 12660012), 1x MEM Non-Essential Amino Acids (Thermo Fisher, Cat. No. 11140050), 1x N-2 Supplement (Thermo Fisher, Cat. No. 17502048), 10 ng/mL NT-3 (PeproTech, Cat. No. 450-03), 10 ng/mL BDNF (PeproTech, Cat. No. 450-02), 1 µg/mL mouse Laminin (Thermo Fisher, Cat. No. 23017015), 10 nM ROCK inhibitor, and 2 µg/mL doxycycline hyclate (Sigma-Aldrich, Cat. No. D9891) to induce *NGN2* (**Fig. S3**). 60 million selected iPSCs were plated across three 15cm dishes (20 million cells per dish in 30 mL predifferentiation medium each). Media was replaced on days −2 and −1 with the same formulation lacking ROCK inhibitor. On day −1, three 15cm dishes and four wells of a 48 well plate were coated with 15 µg/mL Poly-L-Ornithine (Sigma-Aldrich, Cat. No. P3655) in DPBS, incubated overnight at 37°C, then washed three times with DPBS and dried in a 37°C incubator. On day 0, pre-differentiated cells were dissociated and replated in maturation medium composed of 50% Neurobasal-A (Thermo Fisher, Cat. No. 10888022) and 50% DMEM/F-12 (Thermo Fisher, Cat. No. 11320033), supplemented with 1x MEM Non-Essential Amino Acids, 0.5x GlutaMAX (Thermo Fisher, Cat. No. 35050061), 0.5x N-2, 0.5x B-27 (Thermo Fisher, Cat. No. 17504044), 10 ng/mL NT-3, 10 ng/mL BDNF, 1 µg/mL Laminin, 2 µg/mL doxycycline, and 20 µM trimethoprim (TMP; Sigma-Aldrich, Cat. No. 92131) to stabilize and activate the ecDHFR-tagged CRISPRa system (**Fig. S3**). 120 million cells were plated across the three 15cm dishes (40 million cells per dish in 40ml of maturation medium each). 100,000 neurons were plated in 400ul of maturation medium in each of 4 coated wells of the 48 well plate. On day 7, half the medium was replaced with fresh maturation medium lacking doxycycline; on day 14, half was replaced with twice that volume of doxycycline-free medium. Cells in the 48 well plate were used to confirm functional CRISPRa machinery using a minP-tdTomato reporter assay^59^ (**Fig. S3**) as previously described^22,59^. Neurons were harvested for single cell RNA-seq on day 19.

#### Cerebral organoids

HUES66-derived *CHD8* and *SCN2A* mutant stem cells were established as previously described^97,98^. Human cortical organoids were produced as previously published^65,66^. Briefly, stem cells constitutively expressing dCas9-p300 (*pLV-dCas9-p300-P2A-PuroR* was a gift from Charles Gersbach (Addgene plasmid # 83889; http://n2t.net/addgene:83889 ; RRID:Addgene_83889)) were seeded into an Aggrewell 800 tissue culture plate (Stem Cell Technologies 34815) at 3*10^6 cells per well in 1.5 mL E8 media (Thermo). Every 24 hours for 3 days, media was replaced without disrupting the embryoid bodies in the microwells with E6 media containing 5uM dorsomorphin (PubChem, Ref. No. 11524144), 10 uM SB431542 (PubChem, Ref. No. 4521392), and 2.5 uM XAV939 (PubChem, Ref. No. 329830802). On the fourth day, embryoid bodies were collected from each well with a wide bore pipette tip and a 40 um strainer (Pluriselect, Cat. No. 43-50040-51) to be plated in 10 cm ultralow attachment tissue culture dishes (Corning, Cat. No. 4615; Sbio, Cat. No. MS-90900Z). The culture was undisturbed for the fifth day. After an additional 2 days of culture in E6 media with dorsomorphin, SB431542, and XAV939 with daily changes, the media is changed to Neurobasal A (ThermoFisher Scientific, Cat. No. 10888022) supplemented with B27 without vitamin A (ThermoFisher Scientific, Cat. No. 12587010), GlutaMAX (ThermoFisher Scientific, Cat. No. 35050061), 20 ng/ml EGF, and 20 ng/ml FGF and replaced daily until day 17 post-initiation. From day 18-25, media was replaced every other day. From day 26-43, the media was replaced every other day with Neurobasal A with B27 without vitamin A, GlutaMAX, 20 ng/ml BDNF, and 20 ng/ml NT3. After day 43, media is replaced approximately every four days with Neurobasal A with B27 without vitamin A and GlutaMAX.

#### Patient iPSC lines

Patient iPSC lines from Cri-du-Chat (Cri1377, Cri1378) and Pitt-Hopkins (PH1, PH2) were obtained from the laboratories of Wendy Chung and Alysson Muotri^27^ respectively. Cells were maintained in StemFlex medium (ThermoFisher Scientific, Cat. No. A3349401) and underwent daily media changes. Cells were routinely passaged following dissociation with accutase (ThermoFisher Scientific, A11105-01) and replated on matrigel (Corning, Cat. No. 356231) coated plates. Media was supplemented with 10 µM ROCK inhibitor Y-27632 (Tocris, Cat. No. 1254) on the day of plating.

#### Primary mouse fibroblasts

Primary mouse fibroblasts from a humanized-Tcf4-Nanoluciferase knock-in mouse were maintained in DMEM (Gibo, Cat. No. 11995065) supplemented with 10% FBS. Fibroblasts were plated at 5,000 cells/well in black-walled 384-well plates for luminescence experiments or at 25,000 cells/well in 24-well plates for qRT-PCR experiments with 1% Pen/Strep (Gibo, Cat. No. 15140122).

### Construct delivery

#### iPSC-derived neuron lentiviral transduction for MPRA

Following ½ media change at DIV7, NGN2 neurons were infected with high MOI lentivirus via magnetotransduction. For titrations, 0, 1, 2, 4, 8, 16, 32, or 64 uL of lentivirus were combined with 6 uL of ViroMag R/L transduction reagent (OZ Biosciences, Cat. No. RL41000) and incubated for 20 min at RT. Mixtures were added dropwise to ∼120k cells per well on a 24 well plate and incubated on a magnetic plate (OZ Biosciences, Cat. No. MF14000) for 20 min. Following magnetotransduction, cells were removed from the magnetic plate and cultured until harvesting at DIV14. For functional titrations, gDNA was harvested using the Wizard SV genomic DNA purification kit (Promega, Cat. No. A2361) and functional titers were calculated via qPCR by comparing Ct values from LP34, WPRE, and BB (**Table S7**). For MPRA, high titers are required to achieve sufficient representation across barcodes and cCREs, 6× 10 cm plates with ∼7.5 M cells per plate (2 plates, 15 M cells per replicate) were transduced with 300 uL of concentrated lentivirus and 150uL of ViroMag R/L per 10cm plate (600 uL total lentivirus per replicate). We achieved a mean MOI of 46 across 3 replicates.

#### Nucleofection, piggyBac transposition, and selection for single cell CRISPRa screening in iPSC-derived neurons

16 million *NGN2-dCas9-VPH* iPSCs (as above) were electroporated with the gRNA library and piggyBac transposase at a 5:1 molar ratio (library:transposase) using P3 reagents and the CB-150 program on the Lonza 4D nucleofector. Cells were distributed across sixteen 100-µL nucleofection cartridges, each loaded with one million cells and 17.5 µg total DNA. Following nucleofection, cells were plated into pre-warmed mTeSR+ basal medium with ROCK inhibitor and maintained at 37 °C. At 24 hours, puromycin selection was initiated at 20 µg/mL (a higher dose was required because the *AAVS1-NGN2* construct already encodes puromycin resistance). Media were changed daily with mTeSR+ basal medium containing ROCK inhibitor and 20 µg/mL puromycin for seven days, after which neuronal differentiation was initiated (see “iPSC-derived neurons for multiplex, single cell CRISPRa screening” under cell lines and culture above).

#### HEK293T

For *SCN2A* and *CHD8* validation experiments, sgRNAs were cloned and packaged into *lentiGuide-Hygro-dTomato* (Addgene, Cat. No. 99376) lentivirus and titered via qPCR. Stably CRISPRa expressing HEK293T cells were plated with ∼200k cells per well on a 12 well plate and cultured for 24 hours prior to infection at MOI=5. Cells were cultured for 3 days following transduction and then harvested with RLT lysis buffer (Qiagen, Cat. No. 74104) supplemented with 0.142M 2-mercaptoethanol (BioRad, Cat. No. 1610710).

For *CTNND2* and *TCF4* validation experiments, sgRNAs were cloned and packed into all-in-one *pLV hU6-gRNA(anti-sense) hUbC-VP64-dCas9-VP64-T2A-GFP* (Addgene, Cat. No. 66707) lentivirus and titered via qPCR. WT HEK293T cells were plated with ∼100k cells per well on a 12 well plate and cultured for 24 hours prior to infection at MOI=5 (for *CTNND2*) or MOI=10 (for *TCF4*). Cells were cultured for 3 days following transduction and then harvested with RLT lysis buffer (Qiagen, Cat. No. 74104) supplemented with 0.142M 2-mercaptoethanol (BioRad, Cat. No. 1610710). Following RNA isolation and cDNA synthesis, strong Sp-dCas9 expression was confirmed for all replicates via qPCR.

#### Patient iPSC line transduction

For *CTNND2* and *TCF4* validation experiments, sgRNAs were cloned and packed into all-in-one *pLV hU6-gRNA(anti-sense) hUbC-VP64-dCas9-VP64-T2A-GFP* (Addgene, Cat. No. 66707) lentivirus and titered via qPCR. Patient iPSCs were plated with ∼100k cells per well on a matrigel coated 12 well plate and cultured for 24 hours prior to infection at MOI=100. Cells were cultured for 3 days following transduction with daily media changes and then harvested with RLT lysis buffer (Qiagen, Cat. No. 74104) supplemented with 0.142M 2-mercaptoethanol (BioRad, Cat. No. 1610710). Following RNA isolation and cDNA synthesis, strong Sp-dCas9 expression was confirmed for all replicates via qPCR.

#### Cerebral organoid transduction

For organoid infection, three organoids per condition were plated in an ultralow attachment 24-well plate (Corning 3473). Lentivirus was infected at MOI 5 for 24 hours. Organoids were collected 1-month post-infection for RNA. For each application, a minimum of three organoids were combined to create one biological replicate for bulk RNA isolation. Organoids were collected and flash frozen in Trizol. RNA isolation was performed using an Arcteus PicoPure RNA Isolation Kit (Thermo KIT0204). cDNA synthesis was performed using a Multiscribe High-Capacity cDNA synthesis kit (Thermo).

#### Primary mouse fibroblasts

Three hours after plating, fibroblasts were dosed with lentivirus encoding dSpCas9-VP64 and either a negative control gRNA or targeting gRNA at MOI=50 and cultured for 7 days.

### Library preparation and sequencing

#### MPRA

Cloned cCREs were matched to their respective 15 bp barcodes via association sequencing. Libraries were amplified from 500ng of MPRA plasmid pool via PCR with NEB Ultra II Q5 mastermix (NEB, Cat. No. M0544S) and P5-pLSmP-ass-i# sample index and P7-pLSmP-ass-gfp custom primers (Cycling conditions: 98°C for 30 seconds, 7 cycles of 98°C × 10 seconds, 65°C x 30 seconds and 65°C x 45 seconds, followed by 65°C x 5 min). The PCR product was purified with 0.6x SPRI select beads (Beckman Coulter, Cat. No. B23318) and sent for custom sequencing on an Illumina NextSeq 550 with a Mid-Output 2 x 150bp kit (R1:146, R2:146, I1:15, I2:10) by Novogene. The following sequencing primers were used to capture cCREs sequences (pLSmP-ass-seq-R1, pLSmP-ass-seq-R2), unique 15 barcodes (pLSmP-ass-seq-ind), and sample index (pLSmP-rand-ind2) (**Table S7**).

Following MPRA, samples were split into two equal volumes per 10 cm plate and were passed through a QIAshredder column (Qiagen, Cat. No. 79656) to fully lyse cells followed by DNA/RNA extractions with the Allprep DNA/RNA Mini Kit (Qiagen, Cat. No. 80204). Following isolation, RNA samples were treated with TURBO DNA-free kit (ThermoFisher Scientific, Cat. No. AM1907) to remove any residual DNA before starting cDNA synthesis with the SuperScript III kit (ThermoFisher Scientific, Cat. No. 12574026). gDNA and cDNA were concentrated to 160 ng/uL via ethanol precipitation prior to sample indexing via PCR. DNA and RNA samples for each replicate were amplified with NEBNext HF 2x master mix (NEB, Cat. No. M0541L) and P7-pLSmP-ass16UMI-gfp and P5-pLSmP-5bc-i# custom primers (Cycling conditions: 98°C for 2 minutes, 3 cycles of 98°C × 10 seconds, 72°C x 15 seconds and 72°C x 20 seconds, followed by 72°C x 2 min). The PCR product was purified with 1.4x SPRI select beads and a second round PCR was performed using NEBNext HF 2x master mix and Illumina P7, P5 primers (Cycling conditions: 98°C for 2 minutes, 12-14 cycles of 98°C × 10 seconds, 72°C x 15 seconds and 72°C x 20 seconds, followed by 72°C x 2 min). Final DNA and RNA barcode libraries were purified with 1.2x SPRI select beads and sent for custom sequencing on an Illumina NextSeq 550 with a NextSeq High Output, 1 x 75 kit (R1:15, R2:15, I1:16, I2:10) by Novogene. The following sequencing primers were used to capture DNA/RNA barcodes (pLSmP-ass-seq-ind1, pLSmP-bc-seq), 16 bp UMIs (pLSmP-UMI-seq), and sample index (pLSmP-5bc-seq-R2) (**Table S7**).

#### Single cell CRISPRa

iPSC-derived neurons were dissociated into single-cell suspensions as described previously^99^. To enrich for cells with higher numbers of integrated gRNAs, dissociated neurons were subjected to fluorescence-activated cell sorting (FACS) for the top 10% of piggyFlex gRNA construct GFP signal immediately prior to single-cell profiling. Cells were captured using 10x Genomics Chromium high-throughput platform (Next GEM Chip M) with the Chromium Next GEM Single Cell 3′ HT v3.1 Dual Index kit and Feature Barcode technology for CRISPR screening (10x Genomics, Document CG000418, Rev C). 16 lanes were processed in total, yielding ∼14,000 neurons per lane.

Final scRNA-seq libraries were first sequenced on an Illumina NextSeq 2000 using a P3 100-cycle kit (R1:28, I1:10, I2:10, R2:90) to confirm quality. Libraries were then sequenced on a NovaSeq6000 S2 flowcell.

### Data preprocessing

#### ABC score

Human promoter regions were obtained from Gencode basic annotation files^42^ filtered for protein coding transcripts and defined as the 500bp or 2000bp upstream of the TSS. Isoform specific promoters with >1bp overlap were merged into a single promoter annotation.

For ABC score analysis, ATAC-seq and H3K27ac ChIP-seq or CUT&RUN data from gestational week 18 (GW18) human prefrontal cortex (PFC)^37^ and 7-8 week NGN2 neurons^35^ were aligned to hg19 using the Encode Consortium ATAC-seq and ChIP-seq pipelines with default settings and pseudo replicate generation turned off^100^. NGN2-iNeuron reads were trimmed to 50bp using TrimGalore with the command --hardtrim5 50 prior to alignment (https://github.com/FelixKrueger/TrimGalore). Output bam files were trimmed, sorted, duplicates and chrM were removed, and merged by biological replicate for ATAC-seq and sorted and duplicates removed for ChIP-seq. Trimmed CUT&RUN reads were aligned to hg19 using Bowtie2 v2.3.5.1 with the following settings --local --very-sensitive-local --no-mixed --no-discordant -I 10 -X 700 and converted to bam files^101^. Duplicated reads were removed from the CUT&RUN bam file using Picard MarkDuplicates v2.26.0 with the -- REMOVE_DUPLICATES =true and --ASSUME_SORTED=true options (http://broadinstitute.github.io/picard/). HiC contacts with 10kb resolution from human GW17-18 fronto-parietal cortex^102^ were filtered for a score > 0 and promoter capture HiC (pcHiC) contacts from 7-8 week old NGN2-iPSC inducible excitatory neurons were obtained from GSE113483^35^. ATAC-seq and H3K27ac ChIP-seq or CUT&RUN bam files were provided as input for ABC score^36^ MakeCandidateRegions.py with the flags --peakExtendFromSummit 250 --nStrongestPeaks 150000. cCREs were provided to the run.neighborhoods.py script in addition to hg19 promoter merged transcript bounds. Finally, predict.py was used to identify gene enhancer contacts using HiC or pcHiC data with the flags --hic_type bedpe --hic_resolution 10000 --scale_hic_using_powerlaw --threshold .02 --make_all_putative.

#### MPRA

Association sequencing data was processed using MPRAflow v2.3.5 association.nf utility with default parameters. DNA/RNA barcode sequencing and coords_to_barcodes.pickle from association sequencing were used as input for the count.nf utility with default parameters.

#### Single cell CRISPRa

Sequencing data were processed with CellRanger v7.0.1. FASTQs were generated with cellranger mkfastq (default parameters), and gene expression matrices with cellranger count against the GRCh38-3.0.0 reference transcriptome (10X Genomics, default parameters). Cells with >20% mitochondrial reads or <900 unique molecular identifiers (UMIs) were excluded. Following quality control filtering, 200,517 neurons were retained. The filtered count matrix was used for all downstream analyses.

### Differential expression analysis

#### MPRA

Downstream MPRA analysis was performed using R v3.4.2 in RStudio v2023.06.1+524. DNA/RNA barcode counts from MPRAflow were combined across sequencing batches and used as input for differential expression analysis with DESeq2^103^ with sequencing batch as a covariate. Active oligos are defined as those with differential expression of RNA vs DNA barcode counts (FDR < 0.01, log2FoldChange > 1). This cutoff corresponds to an empirical FDR of 6.2% based on the count distribution of scrambled negative controls.

#### Single cell CRISPRa

Single cell CRISPR screen analysis was performed using SCEPTRE^22,60,104,105^. Genomic coordinates (hg38) for final gRNA spacers were first isolated using a loop built around the matchPattern() function from the BSgenome package^106^. A global UMI filter of 5 gRNA UMIs/cell was then used to assign gRNAs to single cell transcriptomes^22,31,60^. Count matrices and associated metadata were then used to construct a single cell covariate matrix with gRNA and gene expression library size (total UMIs), unique genes (non-zero expression in gRNA and gene expression libraries), 10X lane, and percent mitochondrial reads input as covariates during model fitting. Genes within a 2 Mb window (1 Mb upstream and downstream) around each gRNA were isolated using the construct_cis_pairs() function. The run_calibration_check() and run_discovery_analysis() functions were used to run one-sided calibration (NTC) and discovery (tests) with default parameters. gRNAs/cCREs were first tested against their predicted target NDD risk genes. Hits from this analysis were then tested against all genes within 1 Mb to assess specificity of upregulations. NTCs were tested against all genes that were tested against targeting gRNAs. Custom R and snakemake^107^ scripts were used to partition tests and parallelize processing. NTCs were randomly downsampled to match the number of targeting *cis* tests for visualization on QQ plots. Resulting SCEPTRE *P*-values were then Benjamini-Hochberg corrected and those < 0.1 were kept for discovery sets.

Single cell CRISPR screen differential expression test *P*-values were visualized alongside tracks for cCREs, target gene links, iPSC-derived neuron ATAC-seq^35^, H3K27ac^35^, and RefSeq validated transcripts (ENSEMBL/NCBI) using the Gviz package^108^.

### Enrichment analysis

For transcription factor motif enrichment analysis, MPRA active oligos were used as input to Homer v5.1^109^, findMotifsGenome.pl with 270 bp window size and inactive oligos as background. Presence or absence of a motif for each cCRE was identified using findMotifsGenome.pl with the -find command for all Homer annotated motifs followed by annotatePeaks.pl for each specific motif instance. All cCREs containing a given motif were averaged by log2(RNA/DNA) count ratio. Biochemical enrichment analysis was performed across 258 ATAC-seq^35^ and histone and transcription factor ChIP-seq^110,111^ datasets. The activity of each biochemical marker was determined using bigWigAverageOverBed from UCSC Genome Browser. Values were averaged for corresponding biochemical marks across datasets and brain regions and correlated with log2(RNA/DNA) count ratios from MPRA. Validated active human enhancers from mouse transgenic assays^55^ were separated by tissue and overlapped with cCREs that contained an oligo in the top 10% of MPRA activity. Odds ratios of mouse transgenic assay activity in neural tissues given the presence of strong MPRA activity were calculated using Fisher’s exact test.

### qPCR

#### HEK293T and patient iPSCs

Samples were harvested and flash frozen with buffer RLT Plus prior to RNA isolation with the RNeasy mini kit (Qiagen, Cat. No. 74104). RNA concentration was measured via Nanodrop and input ng was normalized prior to cDNA synthesis with the SuperScript IV kit (ThermoFisher Scientific, Cat. No. 18091050). Samples were diluted 1:5 in molecular biology grade water and qPCR was performed with SsoFast EvaGreen Supermix (BioRad, Cat. No. 1725204) in a QuantStudio 6 Flex (ThermoFisher Scientific; Cycling conditions: 95°C for 10 min, 40 cycles of 95°C x 15 seconds, 65°C x 1 minute).

#### Cerebral organoids

Quantitative PCR was performed using Luna Universal qPCR Master Mix (New England Biolabs, Cat. No. M3003). Targeting sgRNAs for *SCN2A* and *CHD8* were compared to an empty sgRNA vector co-expressing eGFP (Addgene, Cat. No. 83925).

#### Primary mouse fibroblasts

For qRT-PCR experiments, fibroblasts were washed with PBS, lysed with RLT buffer, and frozen at -80°C until extraction using RNeasy RNA Extraction Kit (Qiagen, Cat. No. 74106). cDNA was synthesized from 200 ng total RNA, and qPCR was performed using PowerUp SYBR Green Master Mix (Applied Biosystems, Cat. No. A25742) on a QuantStudio 5 Real-Time PCR system (Applied Biosystems). Relative expression was analyzed using the comparative CT method and normalized to Eif4a2 expression.

### Luminescence

For luminescence assessment, 25 µL per well of Nano-Glo Luciferase Assay (N1120, Promega) was added to 25 µL of media per well and allowed to incubate for 3 minutes on a rocker at room temperature. Luminescence was measured on a BMG CLARIOstar Plus microplate reader (BMG LABTECH GmbH, Germany). Luminescence values were normalized to that of cells treated with negative control lentivirus.

## Notes

https://github.com/shendurelab/Neural_CRE_MPRA_CRISPRa/

